# Principles of ribosome-associated protein quality control during the synthesis of CFTR

**DOI:** 10.1101/2025.07.11.664365

**Authors:** Tom Joshua Oldfield, Raquel Gonçalves Torres, Romy Enrica Baier, Robin Renn, Débora Broch Trentini

## Abstract

Prolonged translational arrests caused by defective mRNAs activate the ribosome-associated protein quality control (RQC) pathway, which marks harmful incomplete proteins for degradation. Multipass transmembrane proteins have increased propensity to be targeted by the RQC, raising the question of whether problems in transmembrane domain insertion and assembly can also cause RQC-eliciting translational arrests. Here, we investigate RQC-mediated quality control of CFTR, a large transmembrane protein mutated in cystic fibrosis. Reporter assays show that although a fraction of nascent CFTR expressed in HEK293 cells arrests during translation and activates the RQC, multiple interventions compromising CFTR folding and membrane insertion do not exacerbate this response. CFTR translation abortion was also largely unaffected by regulators of translation kinetics such as codon usage, the ribosome collision sensor GCN1, and the SRP ER targeting complex. We propose that the RQC can be triggered by the inherent difficulties in synthesizing transmembrane segments, resulting from their inappropriate interaction with the protein synthesis machinery. Our study uncovers and characterizes a novel physiological role for the RQC in dealing with elongation-arrested transmembrane proteins.

## Introduction

Transmembrane proteins, representing circa 26–36% of the human proteome, mediate critical cellular processes, including cell signaling, adhesion, and transport of a variety of molecules (Fagerberg *et al*, 2010). The synthesis of these proteins entails overcoming various biophysical challenges: aggregation-prone hydrophobic transmembrane segments must be shielded from the crowded aqueous cytosol as well as transported past the polar surface of the membrane into the hydrophobic core of the lipid bilayer. Therefore, the majority of transmembrane proteins, including those of the cell surface and most intracellular compartments, are assembled co-translationally at the endoplasmic reticulum (ER). Accordingly, important milestones in the biogenesis of transmembrane proteins must be achieved concomitantly to their translation: targeting of ribosome-nascent chain complex (RNC) to the ER translocon, establishing correct protein orientation in relation to the membrane, insertion of transmembrane segments (TMs) into the lipid bilayer, and early steps of protein folding and assembly. Moreover, the ER translocon represents a dynamic assembly whose composition is co-translationally tuned to meet the specific requirements of a wide range of substrates. For example, while certain TMs are able to access the membrane via the Sec61 lateral gate, polytopic membrane proteins often require the transient recruitment of alternative insertase complexes such as the ER membrane protein complex (EMC) or the multipass translocon (Chitwood & Hegde, 2020; McGilvray *et al*, 2020; Satoh *et al*, 2015; Shurtleff *et al*, 2018; Smalinskaite *et al*, 2022; Sundaram *et al*, 2022). The presence of such co-translational events imposes temporal constraints on the synthesis of transmembrane proteins, and therefore translation kinetics plays an important role in their biogenesis. For example, many lines of evidence demonstrate that elongation kinetics directly affects the biogenesis of the Cystic Fibrosis Transmembrane Conductance Regulator (CFTR) (Kirchner *et al*, 2017; Oliver *et al*, 2019; Veit *et al*, 2016), a chloride ion channel whose folding is predominantly co-translational (Kim *et al*, 2015; Kleizen *et al*, 2005). Moreover, transcripts encoding transmembrane proteins are prominent targets of GCN1, a ribosome collision sensor that acts as a global regulator of translation dynamics (Müller *et al*, 2023).

While programmed translational pausing represents an important strategy to promote RNC targeting to specific cellular locations, recruitment of interactors, and nascent chain folding (Collart & Weiss, 2020; Komar *et al*, 2024; Stein & Frydman, 2019; Zhang & Ignatova, 2011), prolonged ribosome stalls leading to ribosome collisions may signal for co-translational protein degradation, mRNA decay, and, in more extensive cases, stress response activation (Bengtson & Joazeiro, 2010; Inada & Beckmann, 2024; Juszkiewicz *et al*, 2018; Simms *et al*, 2017; Wu *et al*, 2020). The ribosome-associated quality control (RQC) pathway functions in addressing elongation-stalled polypeptide chains for proteasomal degradation, limiting potentially toxic effects of unfinished proteins that might lack functional domains or that might misfold and aggregate (Bengtson & Joazeiro, 2010; Brandman *et al*, 2012). The ubiquitin ligase ZNF598 (yeast Hel2) is a ribosome collision sensor that marks stalled ribosomes for ribosome splitting, which allows access of the RQC complex to the tRNA-bound nascent chain at the dissociated 60S subunit (Juszkiewicz *et al*., 2018; Juszkiewicz & Hegde, 2017; Shao *et al*, 2013; Sundaramoorthy *et al*, 2017). The core components of the RQC machinery in mammals are the ubiquitin ligase Listerin and the factor NEMF (Bengtson & Joazeiro, 2010; Brandman *et al*., 2012). NEMF recognizes the peptidyl−tRNA molecule at the interface of the dissociated 60S ribosome and recruits Listerin to ubiquitinate the stalled polypeptide near the ribosome exit tunnel, marking it for proteolysis. So far, the RQC has been primarily implicated in the degradation of proteins arising from damaged and defective mRNAs, such as truncated transcripts lacking stop codons or mRNAs carrying a poly(A) tail erroneously placed within the coding sequence. How, and to what extent, cells are able to distinguish between physiological and aberrant ribosome pauses is only beginning to be understood.

Recent studies indicate that the RQC engages in the quality control of complex transmembrane proteins, raising the question of whether the RQC might assume additional functions beyond the surveillance of defective mRNAs (Lakshminarayan *et al*, 2020; Trentini *et al*, 2020). Using MHC-I antigen presentation as a proxy for proteasomal degradation, Trentini *et al*. obtained a dataset of the physiological targets of the RQC ubiquitin ligase Listerin in HeLa cells (Trentini *et al*., 2020). While no overrepresentation for a specific molecular function, cellular component, or biological process was observed in this dataset, the solute carrier (SLC) family of transmembrane transporters stood out, as 5 out of the 100 identified targets belong to this family. Closer inspection of the immunopeptidomics data revealed that the effect of Listerin on protein presentation/degradation increased with the number of transmembrane segments, with proteins containing more than 10 TMs displaying significantly higher RQC-dependent degradation than similarly-sized non-membrane proteins. This tendency was not observed for ER resident, secretory, or monotopic membrane proteins, indicating that TM synthesis rather than ER-localized translation correlates with an increased susceptibility to RQC (Trentini *et al*., 2020).

Translation kinetics can be influenced by multiple factors, including codon usage and charged tRNA availability (Yu *et al*, 2015), unequal rates of peptidyl transfer for different amino acid combinations (Wohlgemuth *et al*, 2008), interactions of the nascent chain with the ribosome exit tunnel (Wilson & Beckmann, 2011), and pulling forces exerted on the nascent chain by co-translational folding, chaperone interactions, and partitioning into the membrane (Goldman *et al*, 2015; Ismail *et al*, 2012; Liu *et al*, 2013; Shalgi *et al*, 2013). We and others previously postulated that failure in co-translational protein insertion and/or assembly at the ER membrane might impact ribosomal elongation, eliciting RQC-mediated co-translational quality control (Eisenack & Trentini, 2022; Lakshminarayan *et al*., 2020; Phillips & Miller, 2020; Trentini *et al*., 2020). In the present study, we use the 12-TM ion channel CFTR as a model case to evaluate the effects of nascent chain folding and membrane insertion on the incidence of terminal ribosome stalling and RQC-mediated degradation of complex transmembrane proteins. While dual fluorescence reporter assays demonstrate that a portion of CFTR translation events in HEK293 cells are prematurely aborted and targeted to RQC-mediated degradation, interfering with CFTR’s ability to fold and assemble in the membrane did not exacerbate RQC activation. Moreover, neither ER stress nor disruption of the SRP ER targeting machinery triggered this response, indicating that RQC does not assume a function in monitoring CFTR folding and assembly. We additionally explored the interplay between the tuning of translation dynamics and the incidence of RQC activation during the synthesis of transmembrane proteins. Strikingly, although the rate of abortive CFTR translation depended on the levels of translation initiation, it was not influenced by regulators of elongation kinetics such as codon usage or GCN1 activity, indicating that the RQC operates at translational arrests that are well distinguished from functional pauses. Taken together, our results suggest that co-translational quality control of CFTR and other multipass membrane proteins can be caused by inappropriate interactions between TM segments and the protein synthesis machinery. We thereby uncover a new function of the RQC pathway in preventing the detrimental consequences brought by the inherent difficulties in synthesizing TM segments.

## Results

### Dual fluorescence reporters reveal that a portion of CFTR translation events result in premature translation abortion and RQC activation

To explore the triggers and mechanisms of co-translational quality control during the synthesis of complex transmembrane proteins, we adapted previously published dual fluorescence reporters (Itakura *et al*, 2016; Juszkiewicz & Hegde, 2017) to quantify RQC-mediated protein degradation and terminal ribosome stalling during the synthesis of CFTR, representing a well-characterized transmembrane protein of medical relevance. To quantify protein degradation, we express GFP and CFTR containing an N-terminal mCherry (mCh) tag from the same mRNA (Figure 1A). These two proteins are separated by a viral 2A peptide sequence, which induces ribosomes to skip the formation of one peptide bond without interrupting ribosome elongation (Donnelly *et al*, 2001). Translation of this construct results in production of the two proteins in equal amount; degradation of the mCh-tagged CFTR causes a reduction of red to green fluorescence ratios as measured by flow cytometry analysis (Figure 1A).

**Figure 1:**
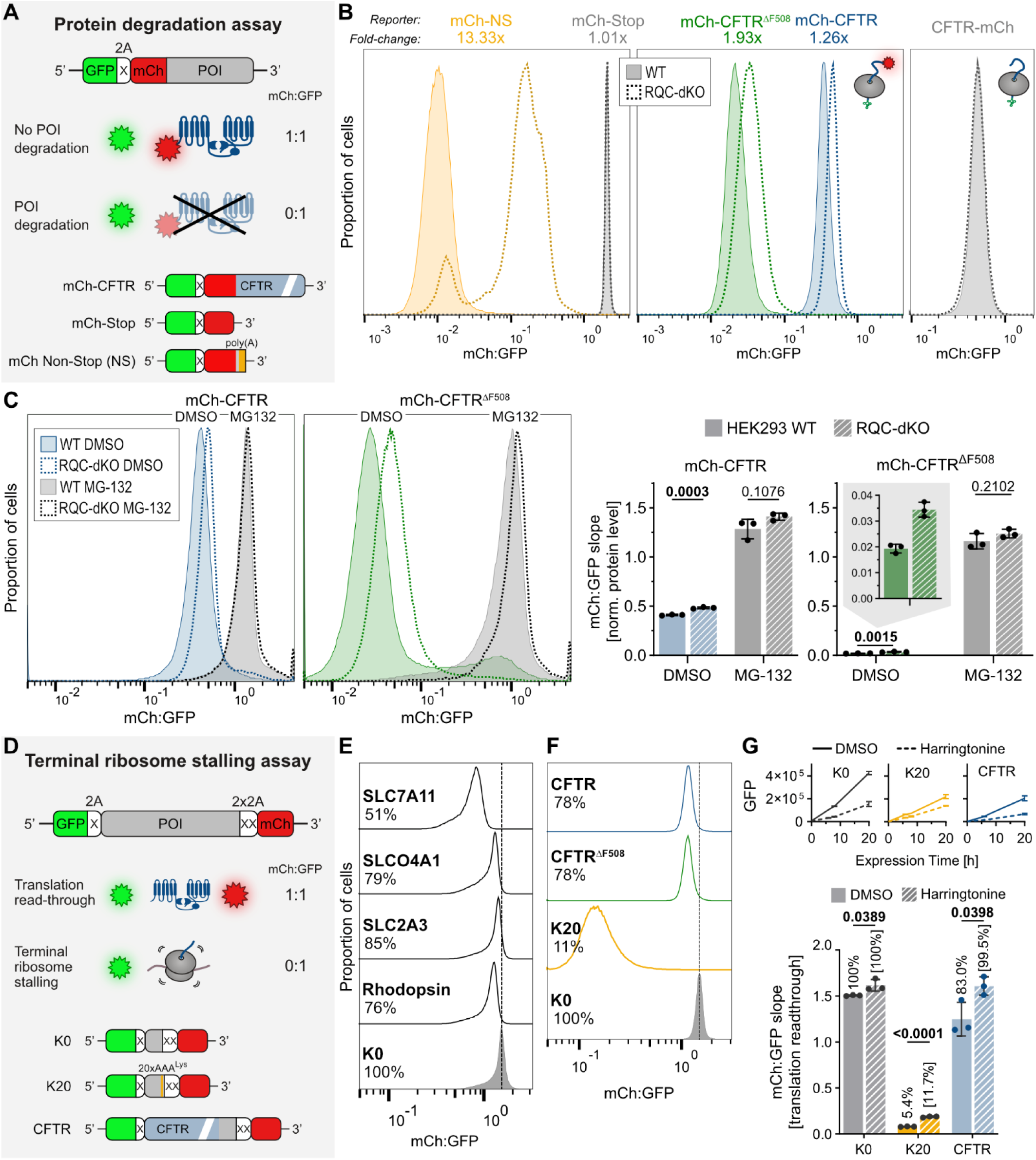
Fluorescence-based reporter assays allow quantification of protein degradation and terminal ribosome stalling. **A** General scheme of protein degradation reporters. Two proteins, GFP and mCh-tagged protein of interest (POI), are produced in equal amounts. Degradation of the POI results in reduced mCh:GFP ratios. **B** Representative histograms of mCh:GFP ratios obtained by flow cytometry analysis of genome-integrated degradation reporters expressed in HEK293 WT or Listerin+NEMF-double KO (RQC-dKO) cells. Fold-changes indicate the dKO/WT ratios of mCh:GFP slopes obtained from Dox time course analyses. **C** Protein degradation analysis of mCh-CFTR and mCh-CFTR^ΔF508^ reporters expressed in the presence of proteasome inhibitor MG-132 in HEK293 WT and RQC-dKO cells. Left: Representative histograms of mCh:GFP ratios obtained by flow cytometry analysis after 24 h of reporter expression under treatment. Right: Mean +/- SD of mCh:GFP slopes of n = 3 independent time course treatments. p = two-tailed unpaired t-tests. **D** General scheme of terminal ribosome stalling reporters. The coding sequence for the POI is placed between GFP and mCh proteins, separated by 2A ribosome skipping sequences. Complete in-frame translation readthrough of the POI results in equal mCh and GFP synthesis. Problems in ribosome elongation culminating in ribosome splitting (translation abortion) preclude synthesis of mCh, resulting in reduced mCh:GFP ratios. K0: non-stalling control sequence; K20: high-stalling control sequence. **E, F** Representative histograms of mCh:GFP ratios obtained by flow cytometry analysis of HEK293 cells expressing the indicated genome-integrated ribosome stalling reporters. Percentages correspond to mCh:GFP ratios normalized to the K0 non-stalling reference, and indicate the relative frequency of complete readthrough of the tested coding sequences. Also see Figure S3. **G** Terminal ribosome stalling analysis of CFTR expressed in the presence of substoichiometric concentration of translation initiation inhibitor Harringtonine. Mean +/- SD of mCh:GFP slopes obtained from Dox time course analyses (n = 3 independent treatments). Percentages correspond to mCh:GFP slopes normalized to the corresponding K0 treatment. p = two-tailed unpaired t-test. Lower graphs show the effect on reporter expression (mean GFP fluorescence intensity) over time.

Cell fractionation experiments demonstrated that the mCh-CFTR degradation reporter expressed in HEK293 cells was efficiently targeted to the ER and presented the typical CFTR glycosylation pattern (Supplementary Figure S1A). As expected, the mCh:GFP ratio largely decreased upon introduction of the cystic fibrosis mutation ΔF508 (mCh-CFTR^ΔF508^, middle panel Figure 1B), which causes global protein misfolding, and upon treatment of cells with an inhibitor of Hsc/Hsp70 chaperones, while it increased upon treatment with CFTR folding correctors VX-809 (Van Goor *et al*, 2011) and VX-445 (Keating *et al*, 2018) (Figure S1B). Moreover, the mCh:GFP ratios of mutant mCh-CFTR^ΔF508^ and, to a lesser extent, mCh-CFTR reporter increased upon disruption of the ubiquitin ligase gp78, an ER-associated protein degradation (ERAD) component previously implicated in the degradation of misfolded CFTR (Ballar *et al*, 2010; Morito *et al*, 2008; Vij *et al*, 2006) (Figure S1C). In conclusion, the mCh-CFTR degradation reporter is able to recapitulate well-established features of CFTR quality control.

To specifically assess RQC-dependent degradation of CFTR, we compared the mCh:GFP ratios in wild-type (WT) and different RQC knockout (KO) HEK293 cell lines we obtained using CRISPR-Cas technology (Figure S1D). As controls of RQC activity, we created two additional protein degradation reporters (Figure 1A): mCh-NS (non-stop, where all stop codons have been mutated to allow translation into the poly(A) tail, representing a difficult to translate sequence that strongly induces ribosome stalling (Chandrasekaran *et al*, 2019)) and mCh-Stop (mCherry containing a stop codon, a non-stalling control) (Figure 1A). As expected, we observed a large (13-fold) increase in the mCh:GFP ratio of mCh-NS in HEK293 Listerin and NEMF double knockout (RQC-dKO) cells in comparison to WT, while the levels of mCh-Stop remained the same (Figure 1B). The expression-normalized levels of mCh-CFTR and mCh-CFTR^ΔF508^ reporters showed a significant ∼1.3-fold and ∼1.9-fold increase in RQC-dKO cells, respectively (Figure 1B and C). This difference was not observed when the reporters were expressed in the presence of the proteasome inhibitor MG-132 (Figure 1C), demonstrating that the observed effect of RQC disruption in the mCh:GFP measurements reflect a loss in proteasome-mediated protein degradation. The relatively modest effect of RQC disruption, compared to the 3.1- and 59.8-fold increases resulting from MG-132 treatment for mCh-CFTR and mCh-CFTR^ΔF508^, respectively, is consistent with previous studies showing that CFTR folding/maturation constitutes an error-prone process subjected to multiple quality control checkpoints (Farinha & Canato, 2017; Sato *et al*, 1998; Younger *et al*, 2006). It is also consistent with the prevailing notion that RQC responds to failed translation events that are relatively rare for each gene product but widespread throughout the translatome.

Since the core RQC complex acts after stalled ribosome splitting (translation abortion), its disruption results in the accumulation of truncated polypeptides but not of full-length protein species (Juszkiewicz & Hegde, 2017; Sundaramoorthy *et al*., 2017). Consistent with this, analysis of a CFTR-mCh reporter, where mCh is fused to the C-terminus of the protein, revealed the same mCh:GFP ratios in WT and RQC-dKO cells (Figure 1B). This finding demonstrates that the observed increase in mCh:GFP ratios of mCh-CFTR upon RQC-dKO reflects RQC-mediated degradation of elongation-arrested protein species, rather than indirect effects on the activity of post-translational quality control pathways acting on the complete protein. Moreover, the effect on mCh-CFTR and mCh-CFTR^ΔF508^ degradation was higher in cells lacking both Listerin and NEMF than in the single knockouts, hinting that CFTR degradation responds to both Listerin ubiquitin ligase activity and NEMF-mediated C-terminal protein extensions (CAT-tails) that serve a degron function (Sitron & Brandman, 2019; Thrun *et al*, 2021), as also observed for the mCh-NS protein (Figure S1E). Expression of Listerin and/or NEMF in the respective knockout cells restored protein degradation, confirming that these effects are specific to RQC disruption (Figure S1E). We additionally tested the effect of the Ubiquitin-fold modifier 1 (UFM1), an ubiquitin-like modifier implicated in the degradation of ribosome-stalled secretory proteins (Scavone *et al*, 2023; Stephani *et al*, 2020; Wang *et al*, 2020). While the degradation of an ER-targeted version of the mCh-NS reporter (ER-mCh-NS) was diminished to a similar extent in both UFM1-KO and RQC-dKO cells, degradation of mCh-CFTR and mCh-CFTR^ΔF508^ was independent of UFM1 (Figure S2). In sum, our experiments demonstrate that a portion of CFTR expressed in HEK293 cells is constitutively subjected to RQC-mediated protein degradation, in line with the notion that large transmembrane proteins are particularly susceptible to this type of degradation (Trentini *et al*., 2020).

To obtain a relative quantification of terminal ribosome stalling, we employed an alternative dual fluorescence reporter where a test sequence is placed between two fluorescent proteins, GFP and mCh, separated by 2A sequences (Figure 1D). GFP fluorescence serves as an expression marker and the mCh signal indicates complete in-frame translation of the test protein. Therefore, decreased mCh:GFP ratios indicate higher incidence of terminal ribosome stalling, *i.e.* stalling events leading to ribosome rescue (ribosome splitting, the commitment step in translation abortion). As a reference for no and high ribosome stalling, we adapted two published reporters: “K0”, a short linker protein, and “K20”, the same linker protein containing 20 consecutive AAA^Lys^ codons, *i.e*., a stall-inducing poly(A) sequence. Our previous immunopeptidome study identified the transmembrane proteins SLC7A11 (12 TMs, 55 kDa) and SLCO4A1 (12 TMs, 77 kDa) as targets of Listerin-mediated degradation, while a similar protein, SLC2A3 (12 TMs, 54 kDa), was not significantly changed by Listerin-KO (Figure S3A) (Trentini *et al*., 2020). We therefore created ribosome stalling reporters of these three proteins by introducing the corresponding coding sequences upstream of the K0 linker. Indeed, we observed that expression of SLC7A11 and SLCO4A1 reporters in HEK293 cells results in lower mCh:GFP ratios than expression of the SLC2A3 reporter (Figure 1E and Figure S3A). Using K0 as a reference for 100% translation readthrough, we estimated 51% (SLC7A11), 79% (SLCO4A1) and 85% (SLC2A3) translation readthrough of these SLC coding sequences. Analysis of the stalling reporter of Rhodopsin (7 TMs, 39 kDa), representing an alternative family of multipass proteins, indicated 76% translation readthrough (Figure 1E and Figure S3A). These results demonstrate that the ribosome stalling reporters are able to recapitulate the translational problem we previously observed for specific endogenous transcripts of HeLa cells. We therefore applied this methodology to quantify the rate of terminal ribosome stalling of the CFTR coding sequence. The CFTR ribosome stalling reporter demonstrated that only circa 78% of CFTR translation events reach the end of the coding sequence (Figure 1F), consistent with our observation that a portion of CFTR nascent chains are targeted to RQC-mediated quality control (Figure 1B and C).

Of note, we observed marked differences in the distribution of mCh to GFP fluorescence ratios between reporters expressed from transiently transfected and genome-integrated reporters, even when comparing cells of equal GFP intensity (Figure S4A). As lipofection can be stressful to cells and results in high cell to cell variation in mCh:GFP ratios (Figure S4A), in the present study we exclusively analyze stalling reporters integrated into the FRT genome locus of Flp-In T-REx 293 cells. In addition, we observed that mCh:GFP ratios can vary across different ranges of reporter expression (*i.e.* GFP intensity), even for single cells measured within the same sample (Figure S4B). Time course analysis of Doxycycline (Dox)-induced expression of genome-integrated stalling reporters indicated that mCh:GFP ratios decrease over time and only reach a stable value after 4 to 8 hours of expression, depending on the reporter (Figure S4C). However, we observed a linear relationship between the mean GFP and mCh fluorescence intensity across the different time point measurements (R^2^ ≥ 0.98) (Figure S4C). The slope of this linear regression (*i.e.* the rate of change of mCh fluorescence in relation to GFP) closely correlates with mCh:GFP ratios obtained after 24 hours of reporter expression (Figure S4C). Therefore, when assessing ribosome stalling in experimental interventions that cause changes in reporter expression, for example due to altered reporter mRNA synthesis/turnover or altered translation rates, we utilize the mCh:GFP slope as a more robust (relative) quantification of terminal ribosome stalling.

Finally, to test if the abortive translation observed with the CFTR stalling reporter originates from ribosome collisions, we treated cells with low concentrations of Harringtonine, an inhibitor of translation initiation that interferes with ribosome elongation during the first rounds of peptide bond formation (Ingolia *et al*, 2011; Robert *et al*, 2009). Partial inhibition of translation initiation promotes wider spacing between ribosomes along the mRNA, and therefore decreases chances of ribosome collisions (Juszkiewicz *et al*., 2018; Kriachkov *et al*, 2023). In line with this, co-treatment of cells with 50 nM Harringtonine upon reporter induction decreased expression (GFP intensity) of the K20 stalling reporter to 63% of DMSO control and improved translation readthrough of the K20 poly(A) sequence from 5.4% to 11.7% of the K0 reference (Figure 1G). In the case of CFTR, Harringtonine treatment reduced CFTR stalling reporter expression to 32% of DMSO control and completely restored CFTR readthrough, from 83.0% to 99.5% (Figure 1G). In conclusion, a portion (∼17-26%) of CFTR translation events in HEK293 cells terminate prematurely, most likely due to ribosome collisions, and elicit co-translational quality control mediated by the core RQC components Listerin and NEMF.

### Problems in folding and membrane insertion do not exacerbate the incidence of abortive CFTR translation

To interrogate whether failure in co-translational CFTR folding and membrane insertion might elicit ribosome stalling and splitting (translation abortion), which is a pre-requisite for RQC recruitment, we tested the effect of a number of interventions that modulate early steps of CFTR folding and/or membrane insertion. First, we determined the effect of the cystic fibrosis mutation ΔF508 on the incidence of terminal ribosome stalling. This mutation affects co-translational folding of nucleotide-binding domain 1 (NBD1) of CFTR, ensuing loss of interdomain interactions and global changes in CFTR conformation (Im *et al*, 2023). Flow cytometry analysis of the CFTR^ΔF508^ stalling reporter revealed an identical rate of translation readthrough as wild-type CFTR (Figure 1F). Treatment of cells with the CFTR folding correctors VX-809, which acts at the early stages of protein synthesis (Kleizen *et al*, 2021), and VX-445, whose folding rescue potential depends on translational dynamics (Kim *et al*, 2023), also had no or only minimal (0.95-fold) effect on the incidence of abortive CFTR translation (Figure 2A), even though these treatments improved CFTR biogenesis (Figure S1B). In addition, treatment of cells with the VER-155008 inhibitor of the Hsc70/Hsp70 chaperones, which aid co-translational CFTR folding (Matsumura *et al*, 2011), markedly decreased the levels of the mCh-CFTR protein in both WT and RQC-dKO cells (Figure S1B and S5B) but had no effect on translation readthrough of the CFTR ribosome stalling reporter (Figure 2A).

**Figure 2:**
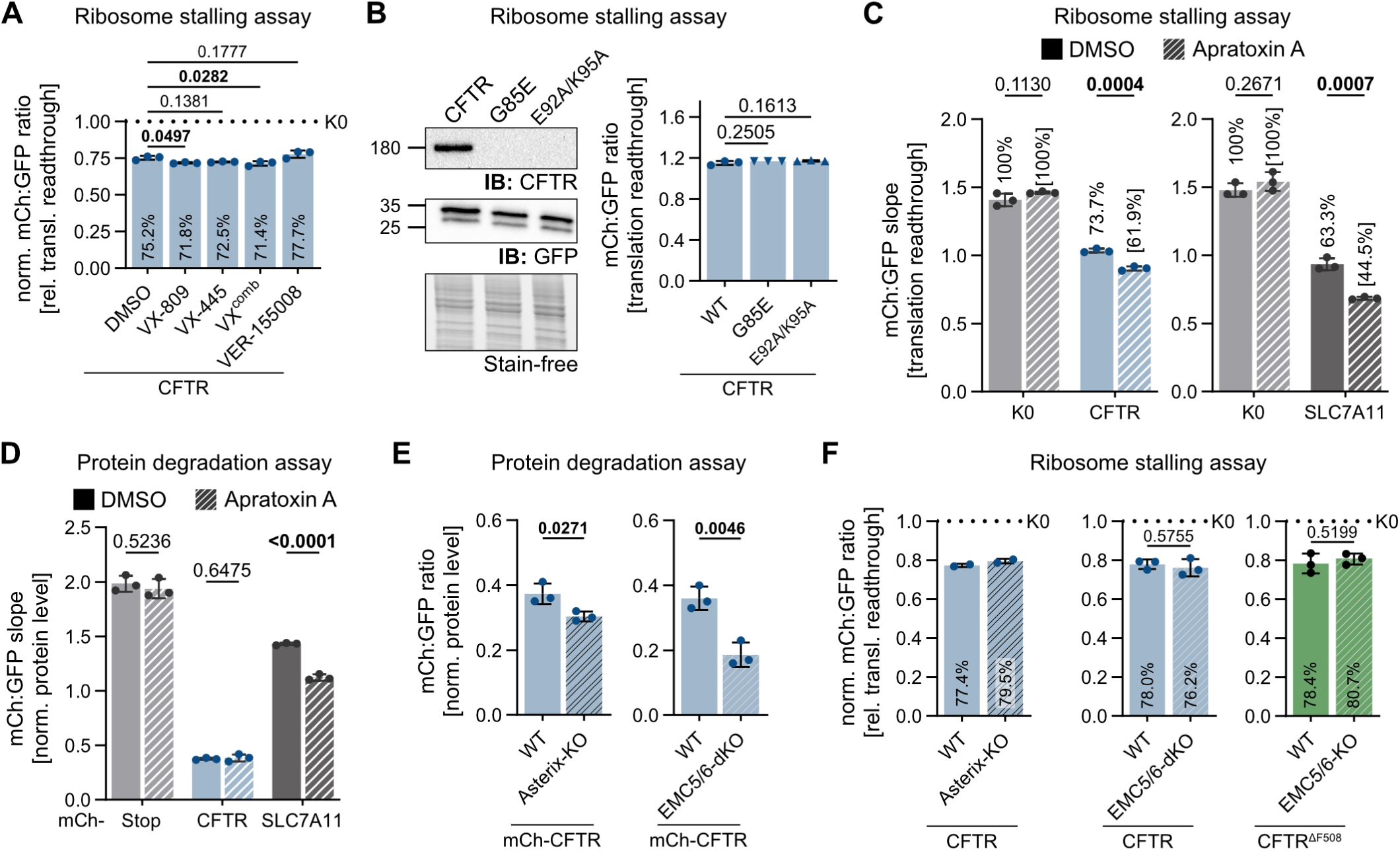
Effect of interventions in protein folding and membrane insertion on translation abortion of CFTR. **A** Terminal ribosome stalling analysis of CFTR expressed in the presence of folding correctors (VX-809, VX-445, or a combination of both) or Hsp70 inhibitor (VER-155008). Graph depicts mean +/- SD of mCh:GFP ratio of the CFTR stalling reporter normalized to the respective K0 control (n = 3 independent treatments). Only cells presenting the same expression range across different treatments (GFP intensity gate, Figure S5A) were included in the analysis. p = one-way ANOVA with Dunnet’s correction. **B** Terminal ribosome stalling analysis of CFTR containing signal anchor mutations. Left: Immunoblot analysis of CFTR variants expressed from the stalling reporters. Right: Mean +/- SD mCh:GFP ratio of genome-integrated CFTR stalling reporters (n = 3 independent analyses of each polyclonal reporter cell line). p = one-way ANOVA with Dunnet’s correction. **C, D** Terminal ribosome stalling (C) and protein degradation (D) analyses of CFTR and SLC7A11 reporters expressed in the presence of Sec61 inhibitor Apratoxin A. Graph displays mean +/- SD of mCh:GFP slopes obtained from time course analyses (n = 3 independent treatments). Percentages indicate relative translation readthrough compared to the corresponding K0 treatment. p = two-tailed unpaired t-tests. **E, F** Protein degradation (E) and terminal ribosome stalling (F) analyses of CFTR reporters expressed in WT, Asterix-KO and EMC5/6-dKO HEK293 cells for 24 h. n = 2 or 3 independent analyses of each polyclonal reporter cell line. p = two-tailed unpaired t-tests.

Next, we addressed the effect of interfering with CFTR targeting and insertion into the ER membrane. To that end, we created two CFTR ribosome stalling reporters containing mutations in its signal anchor (TM1) sequence. According to published *in vitro* translation experiments, G85E is a cystic fibrosis mutation that decreases TM1 hydrophobicity and compromises its ability to direct protein insertion into the ER; and the combined E92A/K95A mutations increase TM1 hydrophobicity and therefore improve signal sequence activity (Lu *et al*, 1998), although other TMs also contribute to the targeting of CFTR to the ER (Carveth *et al*, 2002; Lu *et al*., 1998). Immunoblot analysis indicated that both CFTR^G85E^ and CFTR^E92A/K95A^ protein levels are decreased in relation to wild-type CFTR, demonstrating that these mutations induce a biogenesis defect and consequent protein degradation (Figure 2B). However, both CFTR^G85E^ and CFTR^E92A/K95A^ stalling reporters presented equal rates of translation readthrough as wild-type CFTR (Figure 2B), further demonstrating that membrane assembly defects do not exacerbate terminal ribosome stalling.

In addition, we tested the effect of Apratoxin A, an inhibitor of the Sec61 translocon that binds to the Sec61 lateral gate and inhibits protein translocation or membrane insertion of a wide range of substrates without disrupting RNC targeting to the Sec61 complex (Paatero *et al*, 2016). Apratoxin A treatment resulted in a small but significant decrease in translation readthrough of CFTR, from 73.7% (DMSO) to 61.9% (p = 0.0004), with no effect on the K0 non-stalling control (Figure 2C and S5C-D). A significant decrease in readthrough was also observed for the stalling reporter of the multipass SLC7A11 protein (Figure 2C and S5D) but not for the cytosolic K20 reporter (Figure S5C), indicating that this effect is specific to translation at the ER. Strikingly, protein degradation assays revealed that Apratoxin A increased mCh-SLC7A11 degradation but had no impact on mCh-CFTR protein levels (Figure 2D and S5E) nor on its glycosylation status (Figure S5F), indicating that the opening of Sec61 is not required for CFTR insertion into the membrane. This finding is consistent with recent studies showing that Sec61 lateral gate activity is dispensable for a large subset of polytopic transmembrane proteins that access the ER membrane via alternative chaperone/insertase complexes such as the PAT, GEL, and BOS complexes (multipass translocon) that use the Sec61 as a structural scaffold (McGilvray *et al*., 2020; Smalinskaite *et al*., 2022; Sundaram *et al*., 2022), or the EMC complex that can cooperate with both the Sec61 complex and the multipass translocon (Chitwood & Hegde, 2020; Page *et al*, 2024). Indeed, disruption of the PAT complex subunit Asterix (Figure S5G), which binds to the nascent chain and re-directs it to the multipass translocon (Smalinskaite *et al*., 2022), significantly decreased mCh-CFTR protein levels (Figure 2E). The same was observed for the disruption of the core EMC subunits EMC5 and EMC6 (Figure 2E, Figure S5G). Despite their involvement in CFTR biogenesis, ribosome stalling assays showed that neither Asterix-KO nor EMC5/6-dKO substantially affected the rate of abortive CFTR translation (Figure 2F). Collectively, the results indicate that interfering with CFTR insertion into the membrane is not sufficient to aggravate the translational problem. Assuming that Apratoxin A treatment did not affect RNC targeting to the ER, as indicated by the lack of an effect on CFTR biogenesis, the results also suggest that blocking Sec61 in a closed conformation can impose a physical roadblock to translation—both for proteins that use (SLC7A11) or do not use (CFTR) the Sec61 lateral gate for entry into the membrane.

### The SRP ER targeting system does not influence the rate of abortive CFTR translation

Co-translational targeting of RNCs to the ER is typically mediated by the signal recognition particle (SRP), which recognizes hydrophobic segments (signal peptides and signal anchors) of nascent polypeptide chains and directs them to ER membrane. Association of SRP with the RNC causes a ribosome elongation arrest that is released upon RNC interaction with the SRP receptor (SR) at the ER surface. This translational arrest increases the window of opportunity for the SRP-RNC complex to interact with the rate-limiting SR, thereby maximizing RNC targeting to the ER (Lakkaraju *et al*, 2008; Mason *et al*, 2000; Rapoport *et al*, 1987; Siegel & Walter, 1985; Walter & Blobel, 1981). To test if SRP-mediated translation arrest plays a role in RQC-mediated co-translational quality control of CFTR, we performed siRNA-mediated knockdown of the signal recognition particle subunit 54 (SRP54) and of the SRP receptor subunit alpha (SRPRA) in cells carrying the CFTR or K0 ribosome stalling reporters (Figure 3A). Immunoblot analyses showed that the GFP-normalized levels of the full-length CFTR protein (expressed from the stalling reporter) decreased to approximately 55% (SRP54) and 72% (SRPRA) of non-targeting controls, demonstrating that depletion of these proteins led to a measurable impact on CFTR folding and/or stability (Figure 3B). However, flow cytometry analyses showed that the rate of translation readthrough of CFTR was not significantly changed by SRP or SR depletion (Figure 3C), demonstrating that SRP-mediated ribosome pausing does not play a role in translational arrest of CFTR.

**Figure 3:**
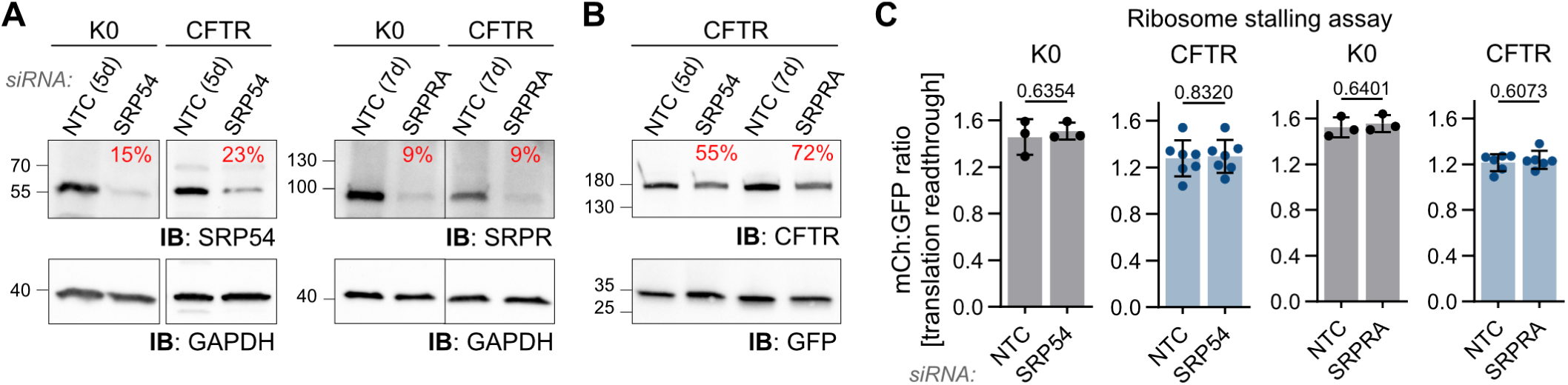
Effect of SRP and SRPR knockdown on translation abortion of CFTR. **A** Immunoblot analysis of SRP54 and SRPR levels upon siRNA-mediated knockdown in cells integrated with K0 or CFTR ribosome stalling reporters. Percentages indicate GAPDH-normalized band intensities in relation to non-targeting control (NTC). **B** Immunoblot analysis of CFTR protein expressed from the ribosome stalling reporter in SRP54 and SRPR knockdown cells. Percentages indicate GFP-normalized CFTR band intensity in relation to the corresponding NTC; reduced percentages indicate protein degradation. **C** Terminal ribosome stalling analysis of CFTR expressed in SRP54 and SRPR knockdown cells. As in A-B, cells were transfected with siRNAs targeting SRP54 or SRPR and analyzed 5 or 7 days post-transfection, respectively. Reporter expression was induced with Dox 24 h prior to harvesting. p = two-tailed unpaired t-test.

Another relevant question is whether the site of translation plays a role in ribosomal stalling during CFTR synthesis. To test if SRP disruption resulted in relocalization of CFTR translation to the cytoplasm, we performed a mild Digitonin treatment to selectively permeabilize the plasma membrane and release soluble cytosolic contents of live cells (Lerner *et al*, 2003; Liu & Fagotto, 2011) (Figure S6A). Quantitative RT-PCR showed that transcripts encoding for the ER resident protein Calnexin (*CANX*) were present at low levels in the cytosolic extract and enriched in the membrane fraction in relation to the mRNA encoding for the cytosolic/nuclear protein Nucleophosmin (*NPM1),* demonstrating efficient separation of cytosolic and ER-bound mRNAs by Digitonin fractionation (Figure S6B). The transcript encoding for the CFTR stalling reporter showed similar levels of cytosol depletion and membrane enrichment as *CANX*, demonstrating that it is efficiently targeted to the ER membrane (Figure S6B). Strikingly, SRP54 or SRPRA knockdown did not significantly increase the fraction of CFTR-or Calnexin-encoding transcripts in the cytoplasmic fraction (Figure S6B). Assuming that mRNA localization correlates with its site of translation, these results indicate that SRP54/SRPRA knockdown did not substantially redirect CFTR translation to free cytosolic ribosomes. Therefore, we cannot exclude that the site of translation influences the rate of translation abortion. While it is possible that in our experimental setup the residual SRP54 and SRPRA levels are sufficient to maintain SRP-mediated targeting activity, previous studies in yeast and human cells indicate that mRNA localization to the ER is largely unaltered by SRP pathway disruption, suggesting the presence of multiple pathways for mRNA association to the ER (Child *et al*, 2023; Kraut-Cohen *et al*, 2013; Pyhtila *et al*, 2008). Nevertheless, taken together with prior *in vitro* evidence implicating the SRP pathway in CFTR membrane insertion (Chen & Zhang, 1996), our finding that SRP54/SRPRA knockdown lowers CFTR protein levels (Figure 3B) supports the conclusion that SRP pathway disruption caused membrane-insertion defects that did not influence translation-abortion rates. In summary, we provide multiple lines of evidence showing that failure in co-translational folding or membrane insertion does not exacerbate premature translation abortion of CFTR, which is required for RQC-mediated protein degradation. A more detailed understanding of the critical determinants of mRNA trafficking and retention on the ER surface will be required to assess how RNC localization affects translation readthrough of CFTR and other transmembrane proteins.

### Proteotoxic ER stress does not promote CFTR translation abortion

Cells respond to proteotoxic ER conditions by activating stress response pathways that mediate the decay of ER-localized mRNAs to acutely decrease secretory/membrane protein synthesis, thereby reducing the burden on the proteostasis network. Both RIDD (regulated IRE1-dependent decay) (Hollien & Weissman, 2006) and ERAS (ER-associated RNA silencing) (Efstathiou *et al*, 2022) pathways promote the degradation of transcripts translated at the ER surface. In principle, such stress response pathways could promote ribosome stalling by cleaving actively-translated mRNAs (with loss of stop codon) or by sterically hindering ribosome movement. To probe the effect of ER stress on co-translational quality control of CFTR, we first inspected whether Dox-controlled overexpression of the CFTR and CFTR^ΔF508^ stalling reporters induces an ER stress response. Immunoblot analysis of two stress markers, the XBP1s and ATF4 transcriptional factors, revealed that in our experimental setup the expression of neither folding-competent nor misfolding-mutant CFTR is sufficient to induce the unfolded protein response (UPR) or the integrated stress response (ISR) (Figure 4A). In addition, treatment of cells with IRE1 endonuclease inhibitor KIRA8 (Harrington *et al*, 2015; Morita *et al*, 2017) did not change the incidence of translation readthrough of CFTR (Figure 4B and S7A). Furthermore, protein degradation assays revealed similar rates of mCh-CFTR^ΔF508^ degradation in HEK293 WT, IRE1-KO (disrupted RIDD), and AGO2-KO (disrupted ERAS) cells (Figure S7B). These findings indicate that the RIDD and ERAS pathways are not responsible for the baseline levels of abortive translation and RQC-mediated degradation we observe with the CFTR dual fluorescence reporters.

**Figure 4:**
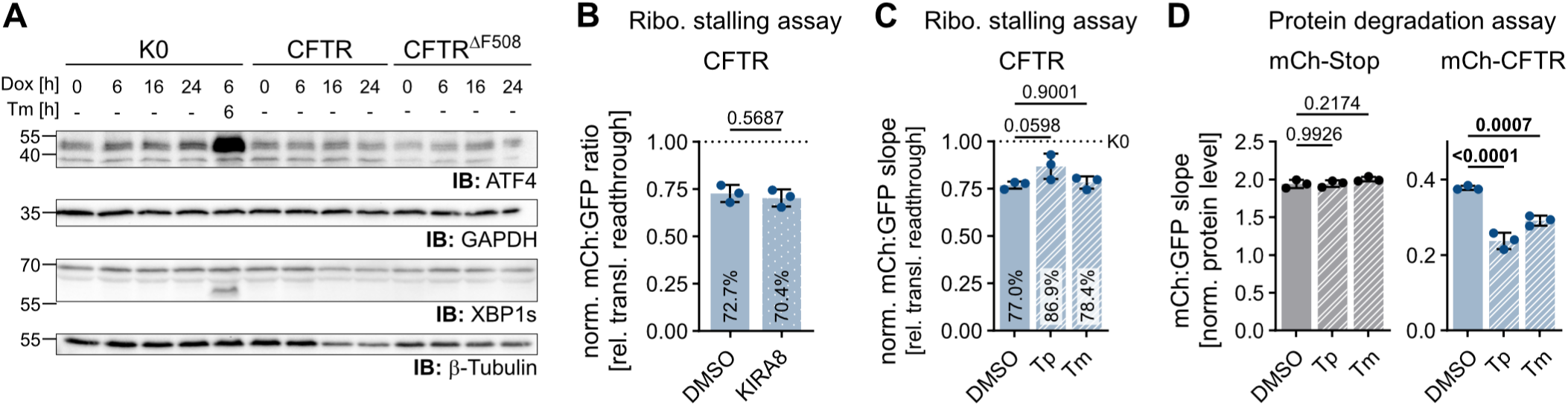
Effect of ER stress on translation abortion of CFTR. **A** Time course analysis of the levels of ER stress markers ATF4 and XBP1s (spliced) upon Dox-induced expression of genome-integrated CFTR, CFTR^ΔF508^, or K0 ribosome stalling reporters. Proteotoxic ER stressor Tunicamycin (Tm) was used as a positive control. **B** Terminal ribosome stalling analysis of CFTR expressed in the presence of the IRE1 inhibitor KIRA8. Mean +/- SD of mCh:GFP ratios of CFTR stalling reporter normalized to the corresponding K0 treatment (n = 3 independent treatments). p = two-tailed unpaired t-test. **C** Terminal ribosome stalling analysis of CFTR expressed in the presence of proteotoxic ER stressors Tunicamycin (Tm) or Thapsigargin (Tp). Mean +/- SD of mCh:GFP slopes normalized to the corresponding K0 treatment (n = 3 independent treatments). p = one-way ANOVA with Dunnet’s correction. See also Figure S7E. **D** Protein degradation analysis of mCh-CFTR and mCh-Stop reporters expressed in the presence of ER stressors, as in C. Mean +/- SD of mCh:GFP slopes (n = 3 independent treatments). p = one-way ANOVA with Dunnet’s correction. See also Figure S7F.

Next, we asked whether ER stress might influence terminal ribosome stalling of CFTR. To that end, we treated cells with proteotoxic ER stressors Tunicamycin (Tm), an inhibitor of N-linked glycosylation that disrupts protein maturation at the ER, or Thapsigargin (Tp), a sarco/endoplasmic reticulum Ca^2+^-ATPase inhibitor that leads to ER Ca^2+^ depletion. Co-treatment of cells upon reporter induction with Dox markedly decreased the expression (GFP intensity) of both K0 and CFTR stalling reporters (Figure S7E), indicating activation of a stress response. Tp but not Tm caused a small non-significant improvement in the readthrough (mCh:GFP slope) of CFTR, from 77% (DMSO) to 87%, using the corresponding treatment of K0 as a reference for 100% readthrough (Figure 4C and S7E). The same treatments resulted in increased degradation of mCh-CFTR protein but not of the cytosolic mCh-Stop control, confirming that the employed ER stressors impact the folding and/or stability of CFTR (Figure 4D and S7F). In conclusion, ER stressors did not increase the incidence of abortive CFTR translation, consistent with our observations that problems in CFTR folding do not induce translation abortion.

However, it is possible that stress-induced attenuation of translation, which might alter the number and the spacing of ribosomes along the mRNA, might mask potential effects of ER stress on ribosome stalling.

### The GCN1 regulator of translation dynamics is not a strong determinant of RQC-mediated quality control of CFTR

To further explore the mechanisms of co-translational quality control during translation of CFTR, we next analyzed the involvement of the ribosome collision sensor GCN1. Müller *et al*. showed that GCN1 senses and stabilizes transient ribosome collisions at slow-decoding suboptimal codons, thereby promoting the interaction of RNCs with co-translational chaperones and the CCR4/NOT mRNA decay complex (Müller *et al*., 2023). The authors propose that GCN1-mediated prolongation of ribosome dwell time may function in increasing the time window for transmembrane protein folding/assembly (Müller *et al*., 2023). The CFTR coding sequence has substantial variation in codon optimality, with suboptimal (slow decoding) codons being particularly frequent in the region downstream of TM6, at a stage where transmembrane domain 1 is likely to assemble (Figure S9A). We therefore hypothesized that the marked susceptibility of CFTR and other multipass proteins towards RQC might represent a trade-off from the tuning of their translation dynamics to meet their specific synthesis requirements. GCN1 also has additional functions in the resolution of certain types of translational roadblocks (Gurzeler *et al*, 2023; Oltion *et al*, 2023; Suryo Rahmanto *et al*, 2023; Zhao *et al*, 2023) (see discussion), and therefore it is also plausible that it might prevent translation abortion.

To assess the interplay between GCN1 and the RQC, we analyzed the effect of GCN1 knockout (Figure S8A) on protein degradation and translation abortion of CFTR and control proteins. Analysis of CFTR and CFTR^ΔF508^ ribosome stalling reporters showed a very small (1.1- and 1.05-fold) decrease in the rate of translation readthrough upon GCN1 disruption (Figure 5A and S8B). Furthermore, protein degradation assays showed that GCN1-KO had no effect on protein levels of mCh-CFTR, but led to an increase (∼1.5-fold) on the levels of the misfolding mCh-CFTR^ΔF508^ mutant (Figure 5B and S8C), which would be consistent with a partial loss of chaperone-assisted protein degradation. The ribosome stalling reporter of SLC7A11, which also contains regions of suboptimal codons (Figure S9A), showed the same levels of translation readthrough in WT and GCN1-KO cells (Figure 5A). We additionally analyzed terminal ribosome stalling of the collagen chain COL4A1, representing a prominent class of GCN1 targets due to their abundance of stall-inducing proline-rich motifs (Müller *et al*., 2023). The results showed no effect of GCN1 on translation readthrough of COL4A1 (Figure 5A), indicating that GCN1 does not signal for translation abortion at problematic sequences. Interestingly, analysis of the K20 stalling reporter revealed a marked (3.7-fold) decrease in translation readthrough of the K20 poly(A) sequence upon disruption of GCN1 (Figure 5A). In agreement with this observation, GCN1-KO further increased the degradation of a short-lived mCh-K20 fusion protein (Figure 5B). These results indicate that during overexpression of a strong inducer of ribosome stalling, GCN1 might play a role in protecting cells from excessive buildup of ribosome collisions and the overwhelming of the RQC system. We deduce that this protective role stems from GCN1’s ability to activate the ISR in situations of high frequency of ribosome collisions, thereby attenuating translation initiation and reducing the incidence of collisions (Müller *et al*., 2023; Nanjaraj Urs *et al*, 2024; Wu *et al*., 2020; Yan & Zaher, 2021; Zhou *et al*, 2025).

**Figure 5:**
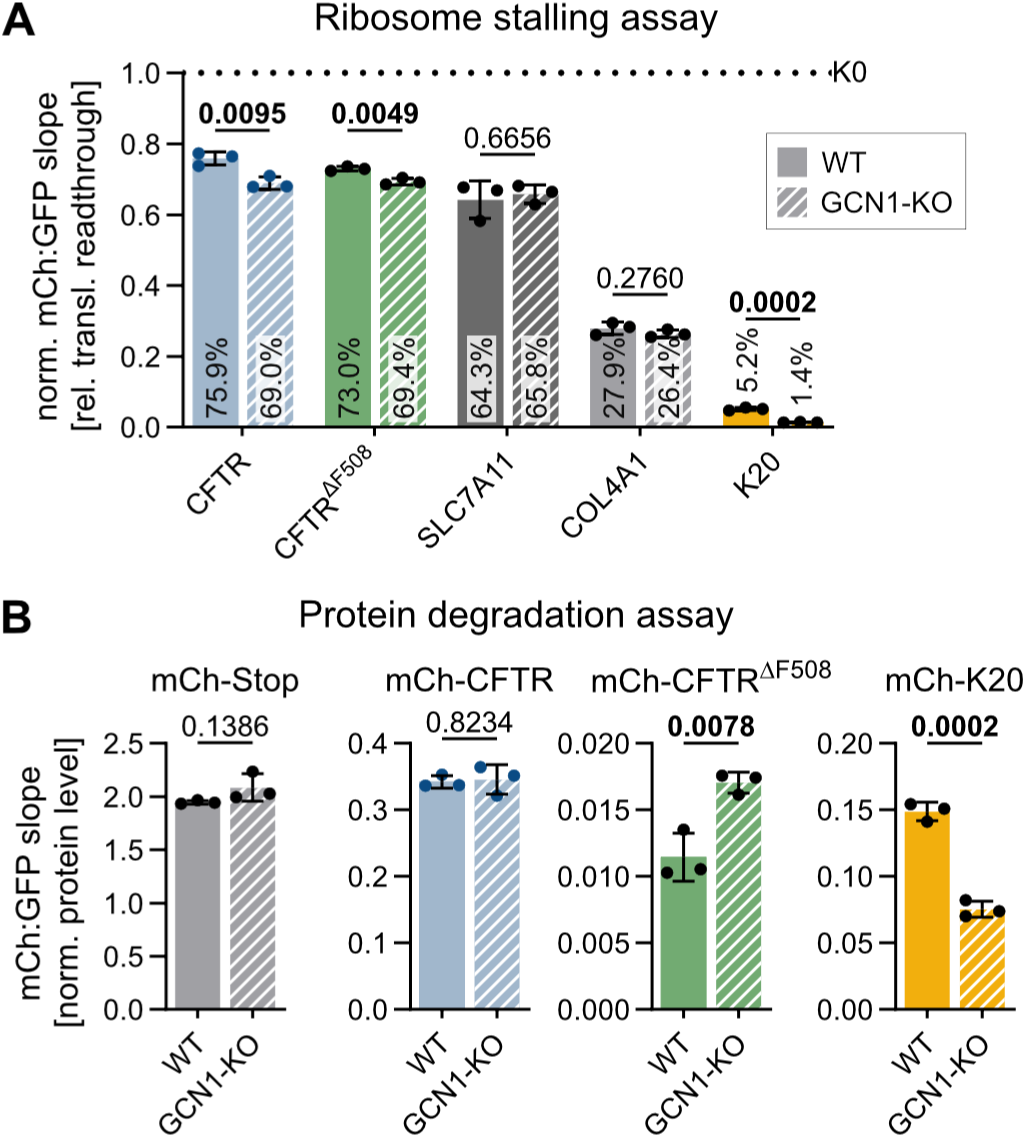
Effect of ribosome collision sensor GCN1 on translation abortion and protein degradation of CFTR and other stalling-prone sequences. **A** Terminal ribosome stalling analysis of CFTR, CFTR^ΔF508^, SLC7A11, COL4A1, and K20 reporters expressed in WT and GCN1-KO HEK293 cells lines. Mean +/- SD of mCh:GFP slopes normalized to K0 expressed in the corresponding genotype. n = 3 independent Dox time course analyses. p = two-tailed unpaired t-tests. See also Figure S8B. **B** Analysis of protein degradation of mCh-Stop, mCh-CFTR, mCh-CFTR^ΔF508^, and mCh-K20 reporters expressed in WT and GCN1-KO cells. Mean +/- SD of mCh:GFP slopes from n = 3 independent Dox time course analyses. p = two-tailed unpaired t-tests. See also Figure S8C.

In summary, our experiments demonstrate that GCN1 does not promote stalled ribosome rescue and RQC activation of the tested examples of stalling-prone transmembrane proteins (CFTR, SLC7A11), excluding this to be the mechanism behind their increased predisposition towards RQC-mediated quality control. Of note, GCN1 disruption presented varied effects on the expression of the different tested reporters (Figure S8B and C), which we presume stems from GCN1’s function in regulating mRNA turnover according to codon optimality (Müller *et al*., 2023). Further studies will be necessary to elucidate whether GCN1 effectively regulates the translation dynamics of these specific proteins.

### Co-translational quality control of CFTR is primarily determined by translation of TM polypeptide segments

Next, we directly assessed how codon usage, a determinant of translation dynamics, influences RQC-mediated quality control of CFTR and SLC7A11. To that end, we created ribosome stalling reporters containing previously published codon-optimized coding sequences where slow-decoding codons were substituted by faster-decoding equivalents (Shah *et al*, 2015; Superti-Furga *et al*, 2020) (Figure S9A). The resulting CFTR^OPT^ and SLC7A11^OPT^ reporters presented markedly increased levels of expression (as measured by GFP intensity) (Figure 6A), consistent with the expected effect of codon optimization on mRNA stability and translation efficiency (Barrington *et al*, 2023; Chu *et al*, 2014; Hia *et al*; Narula *et al*, 2019; Presnyak *et al*, 2015; Wu *et al*, 2019). Strikingly, parallel analysis of native and codon optimized ribosome stalling reporters showed only a minimal (1.03 and 1.11-fold) improvement in translation readthrough (Figure 6A and S9B). This analysis demonstrates that translation abortion and RQC activation during the synthesis of these proteins in HEK293 cells is not caused by stretches of slowly decoded codons, indicating that functional translational pauses are not perceived as a signal for translation abortion. Note that 68% and 63% of the codons of CFTR and SLC7A11 were replaced in their codon-optimized versions, respectively (Figure S10). The observation that such widespread changes in mRNA sequence did not influence the incidence of translation abortion indicates that known sequence-dependent triggers of RQC, such as mRNA secondary structures or near-cognate polyadenylation signals, are not the main cause of the constitutive levels of co-translational quality control of CFTR and SLC7A11 observed in our experiments. It also suggests that their translational problem is not caused by putative sequence-specific RNA binding proteins.

**Figure 6:**
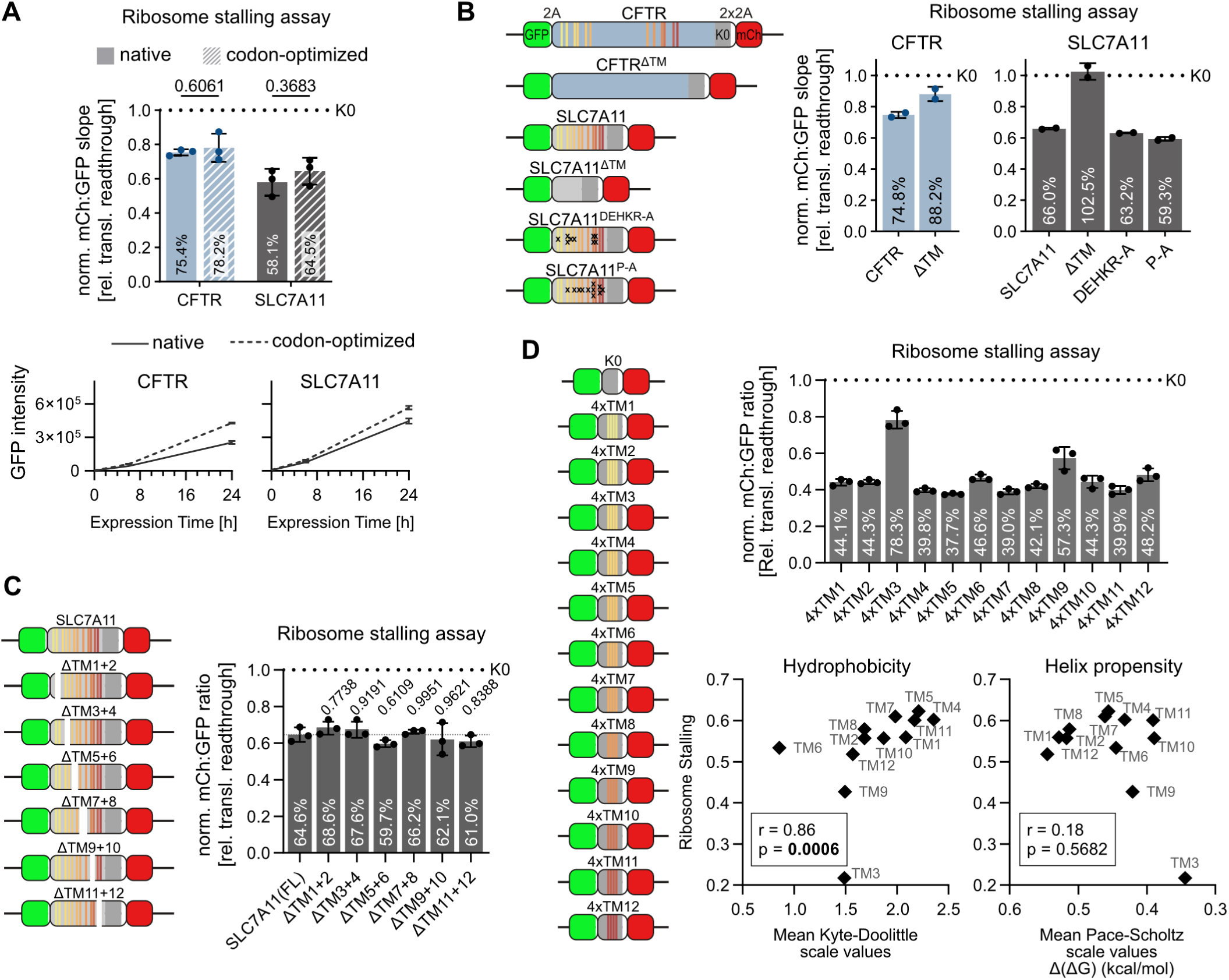
Analysis of the sequence determinants of terminal ribosome stalling of CFTR and SLC7A11. **A** Terminal ribosome stalling analysis of native and codon-optimized CFTR and SLC7A11 coding sequences. Top: Mean +/- SD of mCh:GFP slopes normalized to K0 reference reporter. n = 3 independent time course analyses of each polyclonal reporter cell line, p = two-tailed unpaired t-tests. Bottom: corresponding reporter expression (Mean +/- SD of GFP intensity) upon Dox induction over time. **B** Terminal ribosome stalling analysis of CFTR and SLC7A11 TM mutants. Segments encoding for TMs were removed from the coding sequence (ΔTM). Charged residues (DEHKRA-A) or prolines (P-A) located within TM segments were mutated to alanine (see also Figure S9C). n = 2 independent time course analyses of each polyclonal reporter cell line. **C** Terminal ribosome stalling analysis of SLC7A11 mutants containing the indicated TM pair deletions, including the corresponding inter-TM loop. n = 3 independent time course analyses of each polyclonal reporter cell line. p = one-way ANOVA with Dunnet’s correction, comparison to the full-length (FL) SLC7A11 reporter. See also Figure S9E. **D** Terminal ribosome stalling analysis of K0-derived reporters containing 4 tandem copies of each SLC7A11 TM, separated by a 5 amino acid loop. Mean +/- SD of mCh:GFP ratios normalized to K0 reference reporter. n = 3 independent time course analyses of each polyclonal reporter cell line. Bottom: correlation analysis of terminal ribosomal stalling rate (1-readthrough rate) and the corresponding TM hydrophobicity (mean of Kyte-Doolittle scale values) and alpha-helix propensity (mean Pace-Scholtz scale values, expressed as Δ(ΔG) in kcal/mol relative to Ala). Since low values indicate high helix propensity, y axis is shown in reverse orientation. r = nonparametric Spearman correlation, p = the corresponding p value (n = 12 K0+4xTM reporters). See also Figure S9F.

The results described above support the notion that the translational obstructions leading to ribosome splitting and RQC activation during the synthesis of CFTR and SLC7A11 are caused by the nascent chain itself and not by the coding sequence. As amino acids are added at the peptidyl-transferase center (PTC) of the large ribosomal subunit, the newly synthesized polypeptide chain traverses a long tunnel that can accommodate ∼30-40 amino acid residues. In certain cases, nascent chain interactions with the irregular and chemically complex wall of the ribosome exit tunnel can propagate conformational changes to the PTC that interfere with translation elongation and/or termination (Wilson & Beckmann, 2011). Taking into consideration the unique properties of TM segments, high hydrophobicity and the tendency to adopt α-helical conformation inside the ribosomal tunnel (Bano-Polo *et al*, 2018), we hypothesize that ribosome stalling may originate from improper interactions between TMs and the ribosome. To test this hypothesis, we created CFTR and SLC7A11 stalling reporters in which all TM segments, representing 17% of the CFTR sequence and 48% of the SLC7A11 sequence, were deleted from the coding sequence (ΔTM). Flow cytometry analysis revealed near complete (88%) and complete readthrough of CFTR^ΔTM^ and SLC7A11^ΔTM^ sequences, respectively, demonstrating that translation abortion is indeed largely caused by TM synthesis (Figure 6B).

We additionally created SLC7A11 reporters in which all TM-localized charged amino acids (9 residues, Figure S9C) were substituted to alanine (SLC7A11^DEHKR-A^). The results showed comparable levels (1.04-fold increase) of terminal ribosome stalling between these mutants and wild-type SLC7A11 (Figure 6B and S9D). This indicates that the translational problem is not caused by charge-based interactions between these TM residues and the ribosome exit tunnel. It also supports the notion that poorly hydrophobic TMs that are difficult to partition into the lipid membrane are not the source of the problem. Prolines are often critical residues in many translation arrest sequences due to their poor peptide bond formation kinetics and restricted geometry (Wilson *et al*, 2016). We therefore substituted TM-localized prolines (11 residues) to alanines in the SLC7A11 coding sequence (SLC7A11^P-A^, Figure S9C). However, Pro substitutions did not substantially change the readthrough rate of SLC7A11 (1.11-fold increase; Figure 6B and S9D), indicating that these residues are not essential for this type of ribosomal stalling.

To interrogate which TM segment(s) are responsible for the relatively high rates of terminal ribosome stalling of SLC7A11, we next created a series of SLC7A11 TM deletions. We removed TM segments in pairs, *i.e.* ΔTM1+2, ΔTM3+4, etc, so as to preserve the original membrane orientation of subsequent domains. The resulting ribosome stalling reporters presented very similar rates of translation readthrough as the full-length SLC7A11 coding sequence (Figure 6C and S9E), indicating that terminal ribosome stalling cannot be attributed to specific problematic TM segment(s). To further test this notion, we created ribosome stalling reporters where individual SLC7A11 TMs were inserted into the K0 non-stalling reporter, as 4 tandem copies separated by a 5 residue loop sequence. Flow cytometry analysis showed that all isolated SLC7A11 TMs are capable to induce terminal ribosomal stalling, but to varying degrees (from 21 to 62%, Figure 6D and S9F). Interestingly, we observed a positive correlation between terminal ribosomal stalling and TM hydrophobicity (mean Kyte-Doolittle scale values; nonparametric Pearson r = 0.86) (Figure 6D). However, this was not the case for TM helix propensity (mean Pace-Scholtz scale values) (Figure 6D).

In summary, the SLC7A11-derived reporters demonstrate that the translational problem is not caused by problematic TMs presenting atypical sequence features. Instead, it seems that the inherent characteristics of TMs – namely, their high hydrophobicity – play a role in their inhibitory effect on ribosome elongation. Collectively, our results demonstrate an inherent and complex problem in the synthesis of TM segments, which is likely to contribute to the large dependency of human cells on the RQC pathway for maintenance of proteome homeostasis.

## Discussion

There are several known examples of so-called ribosome arrest peptides, *i.e.* specific amino acid sequences that interact with the ribosome exit tunnel to induce translational stalling as a mean to regulate gene expression, often in response to environmental cues (reviewed in (Ito & Chiba, 2013; Wilson *et al*., 2016)). These translational arrests can be released when the nascent chain experiences pulling forces provided by co-translational folding and/or partitioning into a membrane (Goldman *et al*., 2015; Ismail *et al*., 2012). The previous observation that degradation of transmembrane proteins by the RQC pathway correlates with their number of TM segments (Trentini *et al*., 2020) suggested that these highly hydrophobic sequences may themselves induce translational arrests in human cells. We therefore hypothesized that potentially deleterious translational stalls could arise from interactions of TMs with the ribosome (and/or the membrane insertion machinery) and that, in principle, such interactions could be counteracted by pulling forces provided by productive nascent chain folding and membrane insertion events. Accordingly, activation of RQC would only happen for translation events that failed to achieve certain co-translational milestones. This notion has been corroborated by a study in yeast that showed that problems in membrane insertion by loss of EMC or Sec61 dysfunction led to ribosome stalling and co-translational quality control of certain misfolding mutants of the ABC transporter Yor1 (Lakshminarayan *et al*., 2020). Moreover, selective ribosome profiling experiments in yeast identified Yor1 as one of the many membrane/secretory RNCs targeted by the collision sensor Hel2 when ribosomes failed to associate with the ER (Matsuo & Inada, 2021). In the present study, we provide many lines of evidence demonstrating that this is not the case for human CFTR.

We measured the rate of CFTR translation abortion in the presence of diverse interventions affecting nascent CFTR folding and membrane insertion: signal anchor and misfolding mutations, inhibition of co-translational chaperones, treatment with proteotoxic ER stressors, depletion of ER targeting machinery, and disruption of membrane insertase complexes. Although our experimental setup was sensitive enough to detect terminal ribosome stalling and RQC activation during CFTR synthesis in normal conditions, none of the tested interventions exacerbated RQC-mediated co-translational quality control of CFTR. While characterizing the specific effects of these interventions on the translation kinetics of CFTR is beyond the scope of our study, we can nonetheless conclude that, contrary to our initial hypothesis, the human RQC pathway is unlikely to exert a co-opted function in monitoring the co-translational assembly of CFTR. Of note, although CFTR shares ∼20% sequence homology with Yor1, we demonstrate that CFTR’s membrane insertion pathway is substantially different from the yeast homolog, as it engages with the multipass translocon that may have evolved to increase the efficiency of multipass protein biogenesis in metazoans (Page *et al*., 2024; Sundaram *et al*., 2022). We speculate that the discrepancies observed in the triggers of RQC-mediated quality control during Yor1 and CFTR synthesis may stem from differences in the division of tasks among Sec61, EMC, and other insertase complexes in yeast versus human cells. Moreover, the threshold for ribosome splitting, the commitment step for RQC activation upon ribosome stalling, can be influenced by multiple factors (ribosome density on the mRNA, ribosome modifications, abundance of rescue factors, etc) that are also likely to differ between yeast and humans.

Our observation that a portion of CFTR translation events are aborted midway and consequently targeted to RQC-mediated co-translational degradation is consistent with early studies demonstrating that ribosome-associated nascent CFTR polypeptide chains can be co-translationally modified with ubiquitin (Sato *et al*., 1998), and that the Listerin ubiquitin ligase (a.k.a RNF160) is part of the core CFTR interactome (Pankow *et al*, 2015). Interestingly, a previous study showed that decreasing translation initiation by means of eIF3a silencing, thereby shifting translation of CFTR from polysomes to monosomes, can improve stability, trafficking, and function of multiple disease variants of CFTR expressed in bronchial epithelial cells (Hutt *et al*, 2018). Our results suggest that such effect could be attributed, at least in part, to the prevention of collision-induced translation abortion, as we observed upon Harringtonine treatment.

The in-depth analysis of the CFTR ribosome stalling reporter excludes a number of putative causes for the constitutive levels of abortive translation we measured in our experimental setup, including SRP-mediated translation arrest and the activity of IRE1 and AGO2 endonucleases. The ribosome collision sensor GCN1 has been previously implicated in promoting elongation processivity at problematic peptide motifs (via recruitment of the non-canonical ribosome-binding GTPase 2 – yeast Rbg2/human DRG2) (Pochopien *et al*, 2021), in the resolution of mRNA-protein crosslinks (via activation of the RNF14 E3 ligase) (Suryo Rahmanto *et al*., 2023; Zhao *et al*., 2023), and in the drug-induced degradation of ribosome-trapped eEF1α and eRF1 factors (via activation of the RNF14 and RNF25 E3 ligases) (Gurzeler *et al*., 2023; Oltion *et al*., 2023). The observation that GCN1 knockout had minimal or no effect on terminal ribosome stalling of CFTR and SLC7A11 suggests that their decoding problem is not caused by these types of blockage.

Interestingly, the Sec61 translocon inhibitor Apratoxin A caused an exacerbation of ribosome stalling during both CFTR and SLC7A11 translation, providing a proof-of-principle example that a defective insertion machinery can establish a physical impediment to ribosome elongation. However, considering that CFTR does not depend on the Sec61 lateral gate for entry into the membrane, it is unclear if Apratoxin A promotes inappropriate interactions between the nascent chain and the Sec61 complex itself or if it might (partially) interfere with the recruitment/activity of one or more translocon-associated factors. Our results also demonstrate that it is unlikely that proteotoxic ER stress promotes a similar type of roadblock. Whether, and to what extent, defective membrane insertase complexes might impinge on ribosome elongation in a physiological setting will require a more in-depth understanding of how they interact with the ribosome and the nascent chain itself.

Taking into consideration that ribosomal stalling of the CFTR and SLC7A11 reporters was unaffected by widespread mRNA sequence changes (codon optimized mutants), we propose that the translational problem could be caused by the nascent chain itself, via inappropriate TM-mediated interactions. In principle, these could happen with the ribosome exit tunnel or, alternatively, with membrane insertase complexes, TM-specific co-translational chaperones, or factors involved in RNC targeting to the ER membrane. While the exact interaction(s) leading to an elongation arrest remain to be defined, the observation that disruption of the SRP ER-targeting pathway, the EMC complex, and the multipass translocon did not affect CFTR readthrough levels favors a model where the critical contacts compromising translation elongation happen between TM segments and the ribosome itself. Nevertheless, it should be noted that multiple lines of evidence suggest that ER and cytosol represent distinct biochemical environments for translation, differing in their local abundance of translational regulators, RNA binding proteins, and other protein biogenesis factors (Reid & Nicchitta, 2015). Therefore, even if critical interactions happen with the ribosome, it is possible that the intrinsic property of TM segments in directing translation to the ER surface may also play a role in how they affect ribosome elongation.

Although the terminal ribosome stalling reporters might not recapitulate the exact incidence of translation abortion of native transcripts due to the presence of artificial sequence elements, they provide a comparative measure of the inherent tendency of certain coding sequences towards translation abortion. Our results show that, although a predisposition to abortive translation seems to be a common occurrence among the coding sequences for multipass transmembrane proteins, the incidence of translation abortion can vary widely between different proteins, from 16% (SLC2A3) to 49% (SLC7A11). The exact causes of this variation remain to be fully elucidated. Overall, the transmembrane segments of the tested proteins exhibit similar topology, average hydrophobicity, and amino acid composition (Figure S3B-D). Furthermore, for these selected cases, there was no correlation between coding-sequence length or TM count and the rate of translation abortion, which argues against probabilistic stalling as the primary cause. We hypothesize that the rate of translation abortion measured for the different transmembrane proteins is the result of a complex interplay of multiple factors, including the biochemical properties of TMs, their interactions, the subcellular localization of the RNC, and ribosome density on the mRNA.

In summary, the present study reveals a new class of RQC substrates characterized by the presence of stall-inducing transmembrane protein segments, and characterizes the determinants of RQC activation for this type of substrate. We thereby evaluate emerging concepts of how essential co-translational protein synthesis processes and the tuning of translation dynamics influence RQC activation. We show that while RQC is not exclusively triggered by mRNA defects, it also does not act on the surveillance of problems in protein folding and membrane insertion. We also show that the RQC does not seem to respond to programmed translational pauses that play a physiological role in transmembrane protein biogenesis. Finally, as translation abortion limits the expression of both wild-type and cystic fibrosis-associated CFTR mutants, we envision that preventing translational stalls and/or ribosome splitting could represent a strategy to boost the synthesis of most, if not all, missense CFTR mutants, thereby potentiating the effect of cystic fibrosis drugs acting to improve CFTR folding and activity.

## Methods

### Plasmids and reporter constructs

For the terminal ribosome stalling assay, we adapted the plasmids pmGFP-P2A-K0-P2A-RFP (Addgene plasmid #105686) and pmGFP-P2A-K(AAA)20-P2A-RFP (Addgene plasmid #105688), which were gifts from Ramanujan Hegde. The K0 sequence (147 AA) is composed by 3xFLAG tag, the autonomously folding villin headpiece domain, and the unstructured cytosolic fragment of Sec61β. K20 corresponds to K0 containing a 20xAAA^Lys^ poly(A) stall-inducing sequence. To ensure efficient separation of mCh for all derived constructs, we inserted a T2A site including an 8 amino acid GS linker between the K0 and P2A sequence using Gibson cloning. The reporters were then subcloned into the pcDNA5/FRT/TO vector (Invitrogen). The multiple ribosome stalling reporters introduced in this study were obtained by inserting the protein-coding sequences (CDS) of interest, excluding both initial methionine and stop codon, in frame between the first P2A and the K0 sequence by Gibson cloning. The wild-type CFTR sequence used here corresponds to RefSeq NM_000492.4 containing the rs213950 SNP (MET470VAL natural variant). Site-directed mutagenesis was used to obtain the CFTR^ΔF508^, CFTR^G85E^ and CFTR^E92A/K95A^ mutants. The CFTR^ΔTM^ and SLC7A11^ΔTM^ mutants were designed by deleting all sequences encoding for TM segments according to UniProt annotation. The resulting sequences were ordered as synthetic DNA sequences (GenScript) and introduced to the stalling reporter by Gibson cloning. The SLC7A11^DEHKR-A^ (R48A, R135A, R148A, K166A, K198A, H373A, D384A, D386A, R396A) and SLC7A11^P-A^ (P136A, P208A, P236A, P267A, P320A, P365A, P367A, P374A, P424A, P428A, P462A) TM variants were designed in a similar way, replacing all TM-localized codons encoding for charged residues or prolines to alanine codons, respectively. The coding sequences for SLC7A11 (NM_014331.4) and SLCO4A1 (NM_016354.4) were synthesized by Twist Bioscience. The SLC7A11 stalling reporters containing TM pair deletions (ΔTM1+2, ΔTM3+4, ΔTM5+6, etc) were produced by the plasmid mutagenesis service of the company GenScript. Each deletion comprised the indicated TMs (according to Uniprot annotation) and the corresponding inter-TM loop. Stalling reporters of isolated SLC7A11 TMs correspond to 4 tandem copies of the indicated TM separated by 5 residue spacers (GNFSL). For cloning reasons, each TM and spacer copy corresponds to slightly different codon optimized versions of the same protein sequence, generated with the Codon Optimization Tool (https://eu.idtdna.com/CodonOpt). These sequences were synthesized and inserted at position 85 of the 147 AA residue K0 sequence of the K0 stalling reporter by the cloning service of the company GenScript. Rhodopsin CDS was subcloned from pcDNA3 Rod Opsin, a gift from Robert Lucas (Addgene plasmid #109361). The codon-optimized SLC7A11 sequence was subcloned from pDONR221_SLC7A11, a gift from RESOLUTE Consortium & Giulio Superti-Furga (Addgene plasmid #132244). The SLC2A3 CDS was subcloned from pDONR223_SLC2A3_WT, a gift from Jesse Boehm, William Hahn, and David Root (Addgene plasmid #81787). The COL4A1 sequence was subcloned from cfSGFP2-col4a1, a gift from Ikuo Wada (Addgene plasmid #160317). The SGFP2 and linker sequences contained in the original plasmid were removed during the subcloning process. The codon-optimized (high CAI) CFTR sequence (Shah *et al*., 2015) was a gift from David Müller.

Protein degradation reporters were constructed in the pcDNA3.1(-) or pcDNA5/FRT/TO vector by Gibson cloning based on previously reported constructs (Itakura *et al*., 2016). A synthetic eGFP-P2A-mCh followed by a modified bGH-based UTR sequence lacking stop codons was introduced into pcDNA5/FRT/TO vector to create the mCh Non-Stop (NS) reporter. mCh-Stop corresponds to this construct containing a stop codon at the end of mCh. Protein coding sequences (CFTR, SLC7A11, SOAT1) were introduced after, or in case of the CFTR-mCh reporter, before the mCh coding sequence. ER-mCh-Stop and -NS reporters were created by introduction of the murine MHC-I signal sequence including an N-glycosylation site to the N-terminus of the mCh CDS. The 20xAAA^Lys^ sequence was introduced into the mCh-Stop degradation reporter after the mCh CDS to generate the mCh-K20 reporter.

To avoid toxicity effects of CFTR-containing plasmids in *E. coli* (Gregory *et al*, 1990), for all plasmids containing the full-length CFTR coding sequence the origin of replication was exchanged to a low copy p15A ori obtained from the p15aTet7mut-fuseYFP plasmid, a gift from Lingchong You (Addgene plasmid #28013). Further measures were used to avoid the effects of *CFTR* cytotoxicity in *E. coli*: cloning was typically performed using the Clean Genome *E. coli* strain (Scarab Genomics) and cells were grown at lower temperature (30°C) and higher inoculation densities. Listerin and NEMF expression plasmids were a kind gift from Ulrich Hartl. All reporters were confirmed by sequencing.

### Cell culture

HEK293 (Flp-In T-REx 293, Life Technologies (Invitrogen)) cells were maintained at 37°C, 5% CO_2_ in Dulbecco’s Modified Eagle’s Medium (DMEM) supplemented with 10% FBS (Gibco), 2 mM L-Glutamine (GlutaMAX, Gibco), non-essential amino acids (Gibco), 25 µg/mL Plasmocin (InvivoGen), 10 U/mL Penicillin and 10 µg/mL Streptomycin (Gibco).

DNA transfections were performed using the Lipofectamine 3000 (Invitrogen) reagent according to vendor’s instructions. Typically, 500 ng – 1 µg DNA were transfected with 1.5-2:1 Lipofectamine (volume to mass) ratio in 12-well plates at ∼70% cell confluence. Dox-inducible reporter expression cell lines were generated using the FRT Flp-In system for genome integration according to the vendor’s instructions. Due to technical difficulties, the K0+4xTM9 reporter cell line was not selected with antibiotic after genome integration. Knockout cell lines were revalidated by immunoblotting after reporter integration. All cell lines were routinely tested for contamination with mycoplasma using the qRT-PCR service from Eurofins Genomics. The HEK293 Asterix-KO monoclonal cell line (Chitwood & Hegde, 2020) was a gift from Ramanujan Hegde. The HEK293T gp78-KO monoclonal cell line was a gift from Marius Lemberg and Friederike Korn (Korn *et al*. manuscript in preparation).

### Generation of knockout cell lines

Polyclonal knockout cell lines were generated by employing CRISPR-Cas9-based methods described previously (Lackner *et al*, 2015; Schmid-Burgk *et al*, 2016). Briefly, an sgRNA (see Reagents and Tools Table) sequence targeting an early exon of the protein of interest was designed using the CRISPOR software (Concordet & Haeussler, 2018) and introduced to pSpCas9(BB)-2A-GFP (PX458; Addgene #48138), a gift from Feng Zhang. The gRNA for GCN1 was expressed from a previously published plasmid (Müller *et al*., 2023), a gift from Gopal G. Jayaraj and F. Ulrich Hartl. The sgRNA expression plasmids were co-transfected with one or two additional plasmids: For the CRISPaint method (Schmid-Burgk *et al*., 2016), used for the UFM1-KO, Listerin-KO, NEMF-KO and Listerin+NEMF-double knockout (RQC-dKO) knockout cell lines, a donor plasmid carrying an antibiotic resistance gene and a frame selector plasmid expressing an sgRNA targeting the donor plasmid were used. For the method described by Lackner *et al*. (2015), used for IRE1-KO, AGO2-KO, GCN1-KO, and EMC5+EMC6-double knockout (EMC5/6-dKO) cell lines, a plasmid carrying an antibiotic resistance gene and expressing an sgRNA targeting the flanking regions of the resistance gene was used (Lackner *et al*., 2015). In both cases, the total DNA amount and the ratio between plasmids was determined empirically. Typically, a total amount of 1 µg DNA was transfected per ∼70% confluent well of a 12-well plate. After transfection, cells were allowed to recover at least 48 h before selection with appropriate antibiotic for 2-4 weeks, as needed. The resulting depletion of the targeted protein(s) in the polyclonal KO cell lines was confirmed by immunoblot analysis and fluorescence reporter assays designed to verify previously described knockout phenotypes. For validation of Listerin-, NEMF-, and RQC-dKO cell lines by complementation, protein degradation reporters were transiently co-transfected with Listerin and NEMF expression plasmids in a 1:1 ratio and analysed by flow cytometry as described below.

For functional validation of the IRE1 knockout cell line, XBP1 splicing assays were performed as previously described (Lin *et al*, 2007). In brief, near confluent HEK293 WT and IRE1-KO cells were treated with 1 µM Tp, 2 µg/mL Tm, or 2 µg/mL Tm plus 10 µM IRE1 inhibitor Kira8 for 4 h. Dimethyl sulfoxide (DMSO) served as a solvent control. Cells were harvested by on-plate lysis and total RNA was isolated with the Macherey-Nagel Nucleospin RNA Plus kit following the manufacturers protocol. Total RNA concentration and purity were measured by direct spectrophotometric analysis in a micro-volume spectrophotometer (Nanodrop), and cDNA synthesis was performed with the Applied Biosystems High-Capacity cDNA Reverse Transcription Kit with addition of 1 U/µL murine RNAse inhibitor (New England Biolabs). To amplify the spliced and unspliced *XBP1* mRNA, previously published *XBP1* primers were used (Lin *et al*., 2007), resulting in amplicons of 263 bp for spliced and 289 bp for unspliced *XBP1*. The resulting PCR products were analysed on a 2.5% agarose gel. For functional validation of the AGO2 knockout cell line, AGO2 activity assays measuring the efficiency of siRNA-mediated knockdown of an abundant transcript (*GAPDH*) were performed as described under the “siRNA-mediated knockdown” section and assessed by immunoblot analysis of GAPDH protein levels.

### Dual-color fluorescence reporter assays

For transient expression analysis (Figures S1C, S1E, S2B and S4A), cells were seeded to 12-well plates resulting in a confluency of ∼70% the next day. Reporter plasmids were then transfected using the Lipofectamine 3000 reagent (Invitrogen) according to the vendor’s protocol and analysed by flow cytometry 2 days post transfection. Expression of reporters in pcDNA5/FRT/TO vector was induced by adding Doxycycline (Dox, Sigma-Aldrich) to a final concentration of 1 µg/mL ∼24 h prior to cell harvesting.

For time course analysis, cell lines containing genome-integrated reporters were seeded on 12-well plates and expression was induced by adding Dox to a final concentration of 1 µg/mL for up to 48 h. Chemical treatments were performed at the same time as reporter induction with Dox. Working concentrations for chemicals used in this study are listed in the Reagents and Tools Table. For the duration of MG-132 treatment, media was supplemented with 1 mM L-Cystein and 10 mM L-Asparagine to reduce cytotoxicity of proteasome inhibition (Suraweera *et al*, 2012).

Cells were washed with PBS, detached by trypsinization (TrypLE Express, Gibco) and resuspended in cold medium. The cells were then collected by centrifugation at 4°C at 800 x *g* for 5 min and resuspended in ice-cold PBS supplemented with 3% FBS and 500 ng/mL DAPI. Flow cytometry analysis was performed using a CytoFLEX LX (Beckman Coulter Life Sciences). Data analysis was performed using FlowJo v10 software: Events were gated for (1) average-sized intact cells using the forward vs side scatter, then for (2) single cells using forward scatter peak area vs height. Of those, (3) DAPI-positive events were excluded. For time course analyses, the predominant GFP-positive cell population, *i.e.* cells containing an intact reporter and a functional Dox-induction system, were selected for analysis. Mean GFP and mCh fluorescence intensities of each time point were calculated by FlowJo and imported into GraphPad Prism software. A linear regression between mean GFP and mCh fluorescence intensities was then calculated for each time course replicate. The resulting slopes of these regressions were then evaluated by statistical analysis (two-sided unpaired t-tests for pairwise comparisons and ordinary One-way ANOVA with Dunnet correction for multiple comparisons). For single time point experiments, the Derive Parameters function of FlowJo was used to calculate mCh/GFP intensity ratios of single events, and mean mCh:GFP ratios of the GFP gate were calculated in FlowJo and imported into GraphPad prism for statistical analysis. The GCN1-KO cell line carrying the mCh-CFTR degradation reporter displayed a population which was GFP-positive/mCh-negative with GFP intensity increasing after Dox induction. This population was clearly separated from the main population which responded to Dox with increase in both GFP and mCh intensities. We identified this population as faulty and therefore excluded it from the analysis.

### Estimation of TM hydrophobicity and helix propensity

To estimate TM hydrophobicity, we calculated the mean of the Kyte-Doolittle scale values [from −4.5 (Arg, hydrophylic) to 4.5 (Ile, hydrophobic)](Kyte & Doolittle, 1982) of the residues localized within each TM according to Uniprot annotation. To estimate TM α-helix propensity, we calculated the mean of Pace-Scholtz scale values [expressed as Δ(ΔG) in kcal/mol relative to Ala, 0 = Ala (most favorable helix propensity), 1 = Gly (least favorable helix propensity)] of each TM (Pace & Scholtz, 1998). Proline, representing the worst helix former (substantially below Gly), was not included in this scale because of its atypical characteristics (Pace & Scholtz, 1998).

### siRNA-mediated knockdown

For SRP54 and SRPRA knockdown experiments, semi-confluent 12-well plates of cells containing either K0 or CFTR genome-integrated ribosome stalling reporters were transfected with 25 pmol of Dicer-Substrate siRNA (DsiRNA, IDT) using 2 µL of DharmaFECT1 transfection reagent (Horizon Discovery), following vendor’s recommendations. For SRP54 knockdown, reporter expression was induced with Dox at day 4 post-transfection and harvested at day 5. For SRPRA knockdown, cells were re-transfected at day 4 post-transfection and reporter expression was induced with Dox at day 6 and harvested at day 7. During this time, cells were passaged and/or expanded to 6-well plates as deemed necessary to avoid over-confluence. Experiments were initially performed in 3-4 biological replicates used for flow cytometry analysis and immunoblot validation of protein knockdown efficiency and CFTR protein levels. An additional 3 replicate experiment in the CFTR ribosome stalling reporter cell line was subsequently performed for Digitonin fractionation (qRT-PCR analysis of mRNA localization and mRNA knockdown efficiency) and flow cytometry analysis.

For the AGO2-KO validation assay, HEK293 WT and AGO2-KO were transfected with non-targeting control and GAPDH DsiRNA as described above and harvested at day 3 post-transfection.

### Digitonin extraction of cytoplasmic content

The CFTR ribosome stalling cell line transfected with NTC, SRP54, or SRPRA siRNAs (as described above) were subjected to Digitonin semi-permeabilization with subsequent cellular fractionation to separate soluble cytosolic and membrane-bound fractions, following published protocols (Horste *et al*, 2023; Liu & Fagotto, 2011) with adaptations. Semi-confluent 6-well plates were rinsed once with ice-cold PBS and collected in 500 µL of PBS supplemented with Superase-In RNase inhibitor at 0.1 U/µL (Invitrogen). An aliquot of 90 µL of cell suspension (approximately 18%) was collected as the no extraction control and further diluted with Superase-In supplemented PBS to final volume of 180 µL. An additional aliquot of 30 µL (6%) of cell suspension was used for flow cytometry analysis. The remaining cells were pelleted at 800 x *g* for 4 min at 4°C, resuspended in 180 µl of ice-cold Digitonin solution (80 µg/mL Digitonin, 150 mM NaCl, 20 mM HEPES pH 7.4, 0.2 mM EDTA, 2 mM DTT, 2 mM MgCl_2_, Superase-In 0.1 U/µL), prepared from 1% Digitonin (Ultra-quality, Carl ROTH Art.-Nr. HN76.1) freshly diluted in DMSO. After 10 min incubation on ice, samples were centrifuged again (800 x *g*, 4 min, 4°C) and the supernatant (*i.e.* cytosolic fraction) was collected and the pellet (*i.e.* membrane fraction) was resuspended in 180 µL of PBS supplemented with Superase-In 0.1 U/µL. 100 µg/mL of an RNA co-precipitant (GlycoBlue, Thermo Fisher) was added to all samples and RNA was extracted using 540 µL of TRI Reagent (Sigma-Aldrich) following vendor’s protocol.

### qRT-PCR analysis

Following Digitonin fractionation, purified RNA samples were quantified by direct spectrophotometric analysis in a NanoDrop instrument. For cDNA synthesis, 1 µg of total RNA was reverse transcribed using High-Capacity cDNA Reverse Transcription Kit (Applied Biosystems) according to the manufacturer’s instructions. Resulting cDNA was diluted in nuclease-free water in a 1:10 ratio. Quantitative PCR was performed using NEBNext® Ultra II Q5® Master Mix in a total reaction volume of 10 µL containing 1 µL diluted cDNA template and gene-specific primers (Reagents and Tools Table). The CFTR stalling reporter was detected with primers annealing to the GFP part of the corresponding transcript. Amplification was carried out according to manufactureŕs instructions in a CFX Opus 384 Real Time PCR system (BioRad) and analyzed using the Bio-Rad CFX Maestro 2.3 Software V5.3.022.1030 (BioRad). Abundant transcripts previously described to be highly enriched in the cytosol (*NPM1*) and the ER membrane (*CANX*) (Reid & Nicchitta, 2012) were employed as markers for the corresponding subcellular fractions.

### Subcellular fractionation

For the analysis of subcellular location of the mCh-CFTR and mCh-CFTR^ΔF508^ proteins expressed from the degradation reporter, we employ a cellular fractionation protocol based on differential centrifugation. The same number of cells stably expressing the reporter of interest were seeded to 10 cm dishes. Reporter expression was induced 24 h later by adding Dox to a final concentration of 1 µg/mL. After 48 h of expression, cells were washed with PBS, detached with EDTA-based dissociation buffer (Gibco), collected in medium and washed with ice-cold PBS. Cell pellets were resuspended in hypotonic lysis buffer (10 mM KCl, 1.2 mM MgCl_2_, 10 mM Tris-HCl pH 7.4) supplemented with 1x cOmplete protease inhibitor cocktail (Roche) and 1 mM PMSF. Cells were allowed to swell for 30 min on ice before and then lysed by passing 6 times through a 27-gauge syringe on ice. An aliquot of this total lysate was taken, before nuclei were removed by centrifugation at 1000 x *g*, 4°C for 10 min. The cytosolic fraction and crude membranes were separated by centrifugation at 100,000 x *g*, 4°C for 30 min. The membrane pellet was washed once in hypotonic lysis buffer before resuspending in TNI buffer (Pankow *et al*., 2015) (50 mM Tris pH 7.5, 250 mM NaCl, 1 mM EDTA, 0.5% Igepal CA-630) supplemented with 1x cOmplete protease inhibitor cocktail (Roche), 1 mM PMSF, 2 mM MgCl_2_ and 2 U/mL Benzonase. Igepal CA-630 was added to total lysates and cytosolic fractions to match the concentration in membrane fractions. All samples were sonicated for a total of 2.5 min at 4°C in a waterbath sonicator. Aliquots of all fractions were taken and recombinant PNGase F (purified from *E. coli*) was added to one aliquot of each membrane fractions. All samples were incubated at 37°C for 5 min, then 4x Laemmli sample buffer (Bio-Rad) supplemented with β-mercaptoethanol was added and samples were incubated at 37°C for 20 min before SDS-PAGE and immunoblot analysis as described below.

### SDS-PAGE and immunoblot analysis

Cells were washed with PBS, detached by either trypsinization or with EDTA-based dissociation buffer (when blotting for membrane proteins), collected in medium and washed with ice-cold PBS. Cell pellets were resuspended in either RIPA (50 mM Tris pH 7.5, 150 mM NaCl, 0.5% SDC, 0.1% SDS, 2 mM EDTA, 1% Triton X-100) or TNI buffer (Pankow *et al*., 2015) (50 mM Tris pH 7.5, 250 mM NaCl, 1 mM EDTA, 0.5% Igepal CA-630) each supplemented with 1x cOmplete protease inhibitor cocktail (Roche), 1 mM PMSF, 2 mM MgCl_2_ and 2 U/mL Benzonase and incubated on ice for at least 30 min. Samples were then sonicated for a total of 2.5 min at 4°C in a waterbath sonicator and clarified by centrifugation (17,000 x *g*, 15 min, 4°C). Protein concentrations were determined using the Pierce Rapid Gold BCA protein assay kit (Thermo Scientific). Samples with same amount of total protein were adjusted to the same volume using lysis buffer, then 4x Laemmli sample buffer (Bio-Rad) supplemented with β-mercaptoethanol was added. Samples intended for immunoblotting of membrane proteins were incubated at 37 °C for 20 min, otherwise at 95 °C for 5 min and subjected to SDS-PAGE. When using TGX Stain-Free Precast Gels (Bio-Rad), total protein content was imaged prior to transfer.

For immunoblot analysis, proteins were transferred to 0.45 µm PVDF membranes using the Trans-Blot Turbo Transfer System (Bio-Rad) according to the vendor’s protocol. Membranes were blocked for 1 h at room temperature in 5% milk in TBS-T (20 mM Tris pH 7.5, 150 mM NaCl, 0.1% Tween® 20) before incubation in primary antibody dilution (in 5% BSA supplemented with 0.02% NaN_3_) for 1 h at room temperature or overnight at 4°C. Membranes were washed with TBS-T at room temperature for 3 times of 10 min each after primary and secondary antibody incubations. Signals were detected using HRP-coupled secondary antibodies and Immobilon Classico Western HRP substrate (Millipore). Immunoblot band intensities were obtained using ImageJ (Schneider *et al*, 2012). See Reagents and Tools Table for antibodies used in this study.

## Author contributions

**TJO:** Conceptualization, Data curation, Formal analysis, Investigation, Methodology, Project administration, Validation, Visualization, Writing – original draft, Writing – review & editing. **RGT** and **REB:** Investigation, Methodology, Validation, Writing – review & editing. **RR:** Investigation. **DBT:** Conceptualization, Formal analysis, Methodology, Project administration, Funding acquisition, Supervision, Visualization, Writing – original draft, Writing – review & editing.

## Acknowledgments

This work was funded by the Fritz Thyssen Foundation (project reference 10.22.2.029MN), the Deutsche Forschungsgemeinschaft (DFG) (project number 540625656, CRC1678 project number 520471345), the Center for Molecular Medicine Cologne (JRG-05 and CAP-33), and the Köln Fortune Program/Faculty of Medicine, University of Cologne (project number 376/2021). We thank Uli Kazmaier and Oliver Andler for providing Apratoxin A, David M. Mueller for sharing codon-optimized CFTR plasmids, Ramanujan Hegde for sharing Asterix-KO HEK293 cell line, and Marius Lemberg and Friederike Korn for sharing HEK293T gp78-KO cell line. We thank Gopal Jayaraj for critical reading of the manuscript.

---

**Figure S1:**
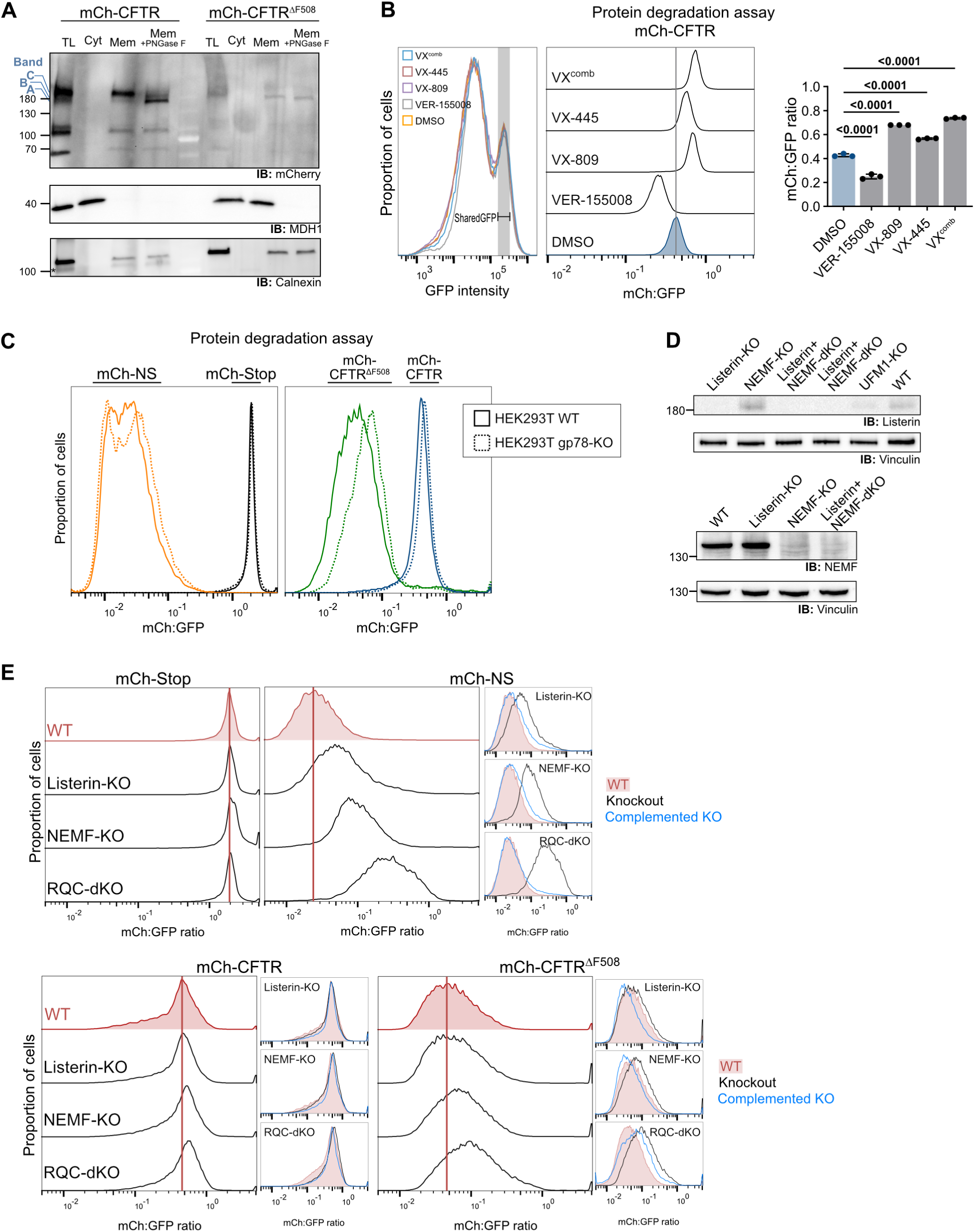
Validation of mCh-CFTR degradation assay reporter. **A** Cellular fractionation analysis of mCh-CFTR and mCh-CFTR^ΔF508^ proteins expressed from genome-integrated protein degradation reporters. Total lysate (TL), cytosolic (Cyt) and membrane fractions (Mem) are shown. Band “B” denotes core glycosylated (ER localized, immature) and “C” denotes fully-glycosylated (mature) mCh-CFTR forms. Band “A” denotes the deglycosylated form obtained by PNGase F treatment. MDH1 and Calnexin serve as controls for loading and enrichment of cytosolic and membrane fractions, respectively. **B** Analysis of mCh-CFTR protein degradation in presence of CFTR folding correctors (VX-809, VX-445, or combined) or Hsp70-Inhibitor VER-155008. To ensure that only cells presenting equal levels of reporter expression across the different conditions are included in the analysis, we employed a GFP gate at an intensity range that is similarly populated across all treatments (SharedGFP, left histogram). Representative histograms of mCh:GFP ratios (middle) and quantification of n = 3 independent treatments (right). p = one-way ANOVA with Dunnet’s correction. **C** Protein degradation analysis in WT and gp78-KO HEK293T cell lines. Cells were transiently transfected with indicated reporters and measured by flow cytometry 48 h later. **D** Immunoblot for validation of Listerin-, NEMF-, and Listerin+NEMF double-KO (RQC-dKO) in HEK293 cell lines. **E** Complementation analysis of protein degradation in WT, Listerin-, NEMF-, and RQC-dKO cell lines. Cells were transiently co-transfected with the indicated degradation reporters and empty vector or expression plasmids for Listerin and/or NEMF. Large histograms (left) show a comparison of mCh:GFP ratios of each reporter across the different knockout cell lines, and small histograms (right) show the effect of Listerin and/or NEMF complementation. Representative histograms from n = 2 independent experiments.

**Figure S2:**
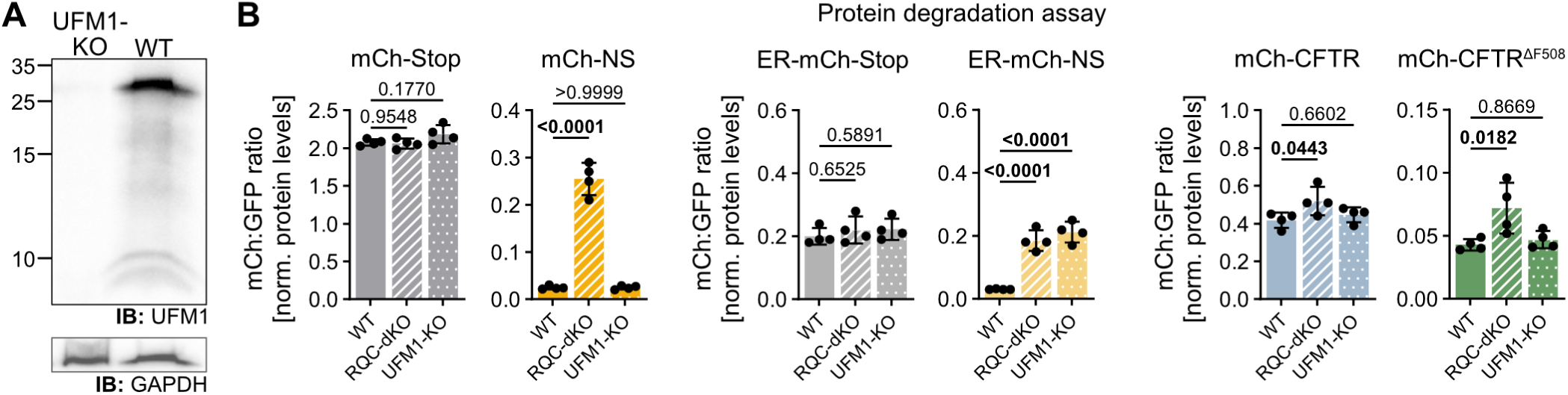
Protein degradation assays in UFM1-KO cells. **A** Immunoblot validation of UFM1-KO in HEK293 cells. **B** Protein degradation analysis in WT, UFM1-, and RQC-dKO HEK293 cell lines. Cells were transiently transfected with the indicated reporters and measured by flow cytometry 48 h later. p = one-way ANOVA with Dunnet’s correction, n = 4 independent experiments.

**Figure S3:**
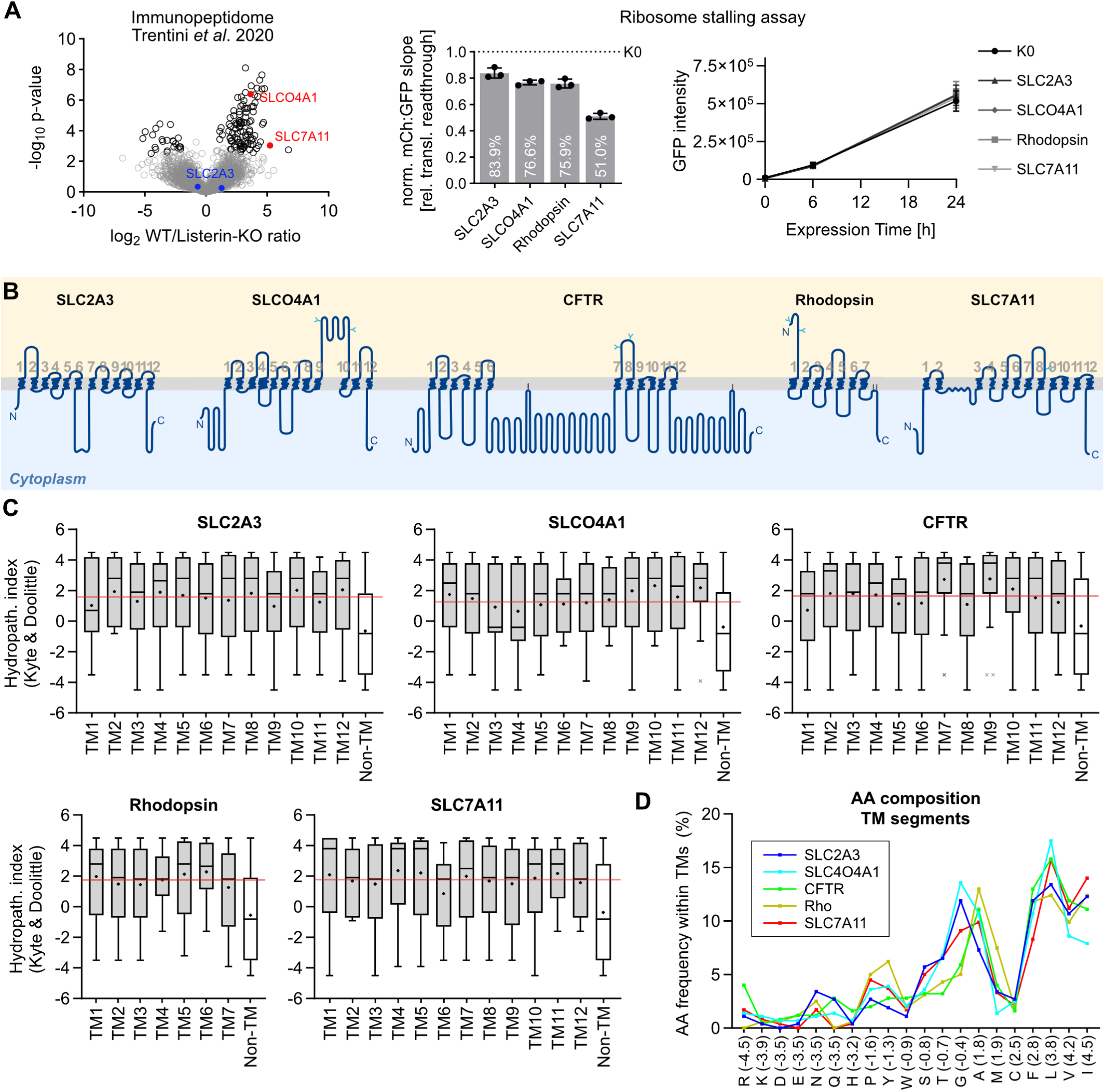
Terminal ribosome stalling assay of selected multipass transmembrane proteins. **A** Left: Volcano plot of previously published immunopeptidome dataset (Trentini *et al*., 2020) comparing MHC-I−bound peptide abundance in WT and Listerin-KO HeLa cells. High WT/KO ratios indicate higher levels of presentation (a proxy for protein degradation) in WT than Listerin-KO cells. Peptides derived from the solute carrier (SLC) transmembrane proteins selected for terminal ribosome stalling analysis are highlighted in red and blue. Center: Mean +/- SD of mCh:GFP slopes of the indicated ribosome stalling reporters normalized to K0 reference reporter. n = 3 independent time course analyses of each polyclonal reporter cell line. p = one-way ANOVA with Dunnet’s correction. Right: corresponding reporter expression (GFP intensity) upon Dox induction over time. **B** Cartoon illustrations of the selected transmembrane proteins. Protein representations are drawn to scale of protein length, according to protein topology and feature annotation derived from UniProt with the help of the Protter interactive protein feature visualization tool (Omasits et al, 2014). Predicted N-linked glycosylation sites are indicated in cyan. **C** Hydrophobicity analysis of the selected transmembrane proteins. Tukey box plot of Kyte-Doolittle scale values for the amino acid residues present in the individual TM segments and Non-TM regions of the indicated proteins. + denotes the mean of the distribution, bar denotes the median. Red horizontal line denotes the mean hydrophobicity of all TM-localized residues of the corresponding protein. **D** Amino acid composition of TM segments of the indicated proteins. X axis shows residues sorted according to the Kyte-Doolittle hydrophobicity scale (values in parenthesis). Y axis shows the relative frequency of each residue within the combined TM segments of the indicated proteins.

**Figure S4:**
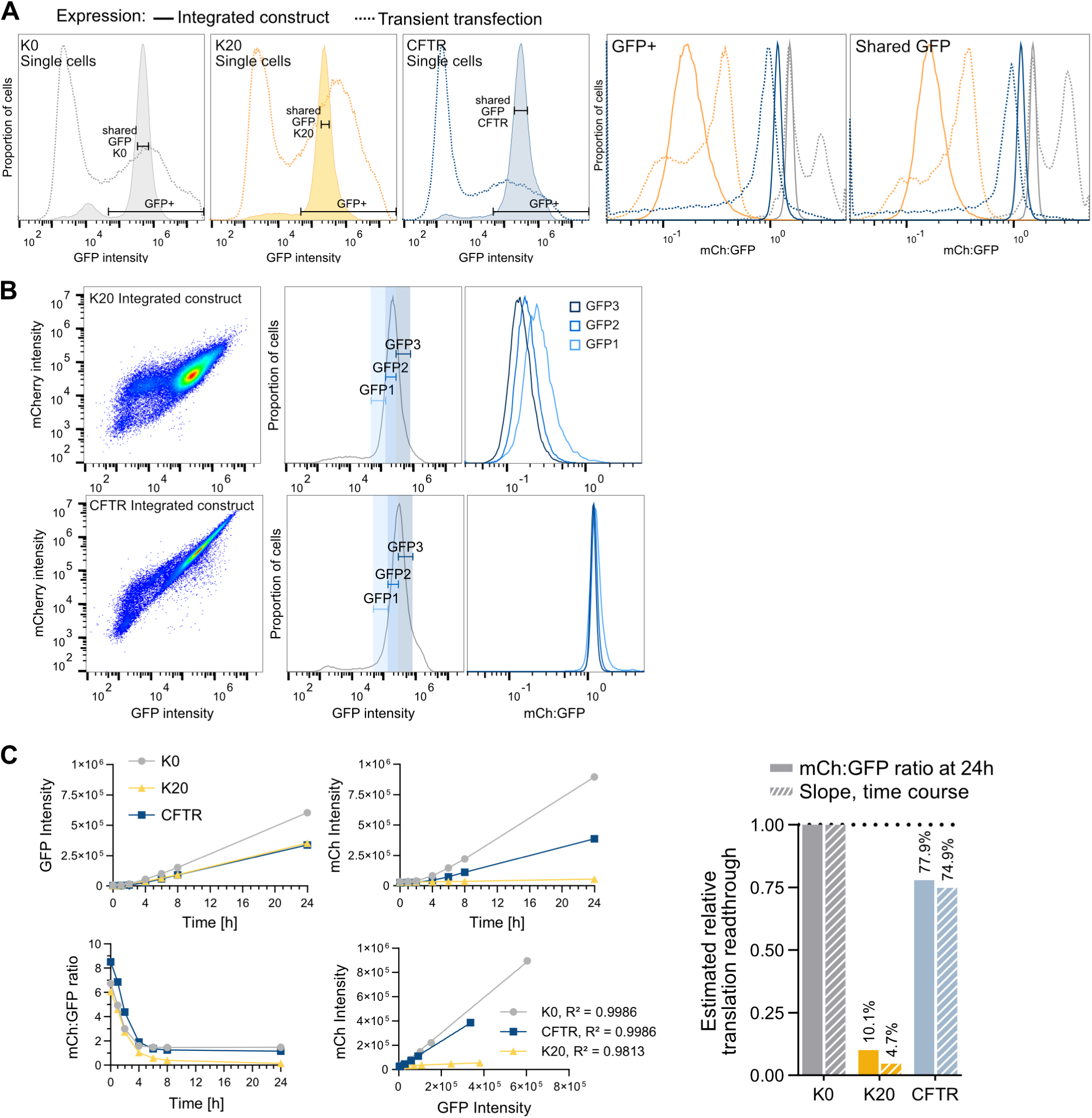
Characterization of terminal ribosome stalling assay. **A** Comparison of terminal ribosome stalling assay results obtained from transiently transfected or genome-integrated reporters. Left: levels of reporter expression (GFP intensity) in each experimental setup. Right: overlay of mCh:GFP ratios of transiently-transfected *vs*. integrated reporters for all GFP+ positive cells and for cells expressing similar levels of reporter (shared GFP gates). **B** Variation of mCh:GFP ratios across different ranges of reporter expression. Left: dot-plots show GFP and mCh fluorescence levels of single cells expressing the K20 (top) and CFTR (bottom) stalling reporters integrated in the same genomic locus. Middle: within the same sample measurement, cells were gated for bottom, average, and top ranges of reporter expression (GFP intensity). Right: mCh:GFP ratios of the depicted reporter expression gates. Despite the variation of reporter expression among isogenic cells being very small, mCh:GFP ratios show considerable variation across the different expression ranges. **C** Time course analysis of CFTR and control ribosome stalling reporters. Cell lines containing genome-integrated Dox-inducible reporters were treated with Dox for 0 to 24 h and analysed by flow cytometry. Line graphs (left) show mean GFP intensity, mCh intensity, and mCh:GFP ratio over time. The equivalent mCh:GFP ratios only reach steady-state levels after ∼6-8 h of expression, depending on the reporter, which precludes using this readout as a measure of terminal ribosome stalling at early time points. However, mean mCh and GFP intensities measured across the different time points follow a linear relationship, with R² indicating high goodness of fit. The slope of the mCh:GFP linear regression quantifies the rate of change of mCh in relation to GFP and therefore can be used as an alternative readout for translation readthrough. Right: relative translation readthrough estimations (K0-normalized values) for mCh:GFP ratio at the 24 h time point and time course slopes.

**Figure S5:**
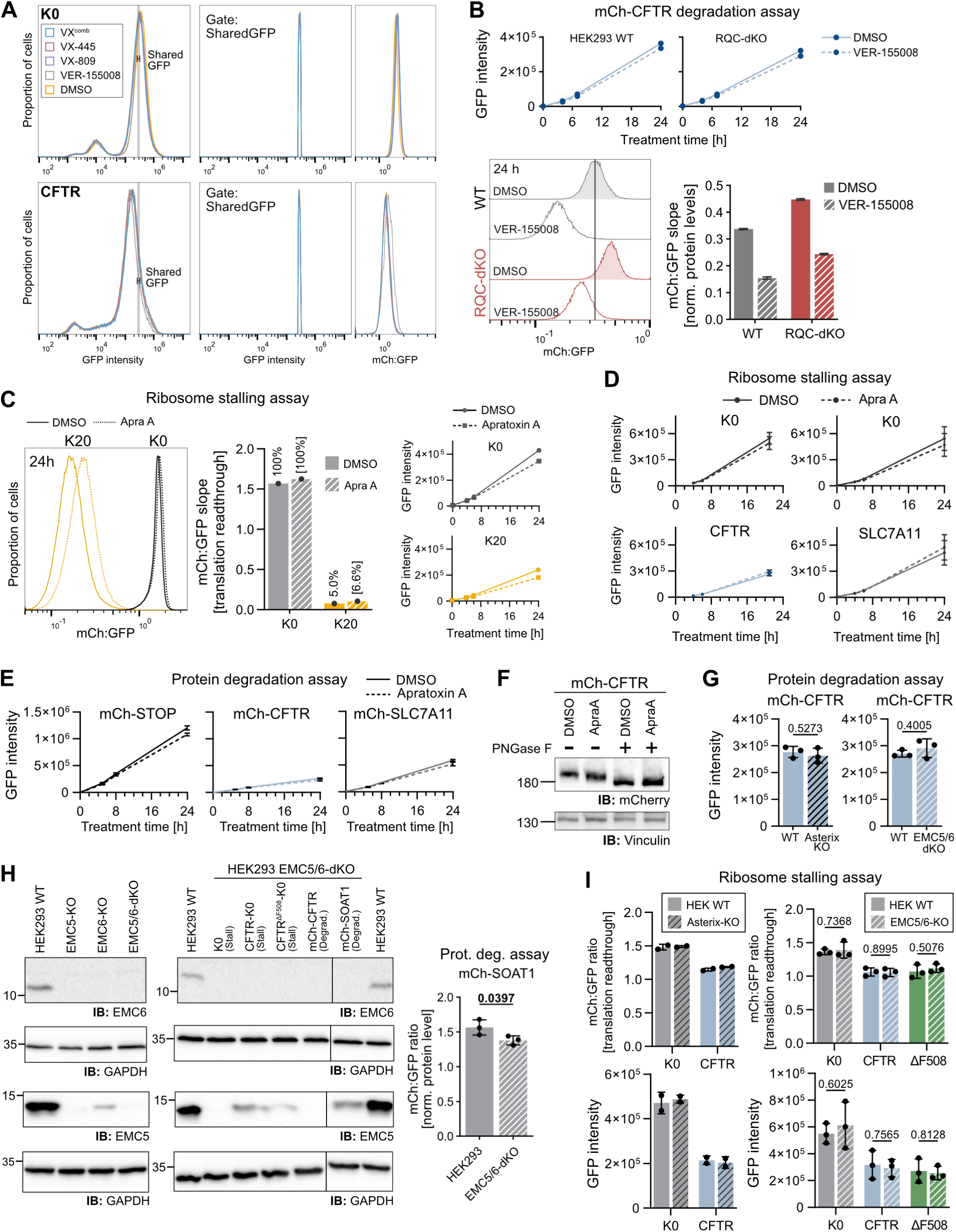
Effect of interventions in protein folding and membrane insertion on translation abortion. **A** Gating strategy for analysis of terminal ribosome stalling of K0 and CFTR expressed in the presence of folding correctors (VX-809, VX-445, or a combination of both) or Hsp70 inhibitor (VER-155008). Left: representative histograms of all measured events. Cells presenting an intermediate expression range populated in both K0 and CFTR reporters (Shared GFP gate) were included in the analysis shown in Figure 2A. Middle and right: representative GFP and mCh:GFP histograms of cells within the Shared GFP gate. **B** Protein degradation analysis of mCh-CFTR in WT and RQC-dKO HEK293 cells in presence of Hsp70 inhibitor (VER-155008) or DMSO control. Top: reporter expression (GFP intensity) upon Dox induction in treated and control conditions over time. Bottom left: histograms of mCh:GFP ratios obtained at the 24 h time point. Bottom right: mCh:GFP slope +/- SEM of linear regression (n = 1) reflect expression-normalized mCh-CFTR protein levels. **C** Terminal ribosome stalling analysis of K0 and K20 control reporters expressed in the presence of Sec61 inhibitor Apratoxin A. Left: mCh:GFP histogram after 24 h of treatment. Center: mCh:GFP slopes obtained from time course analysis (n = 1). Percentages indicate relative translation readthrough compared to the corresponding K0 treatment. Right: reporter expression (GFP intensity) in treated and control conditions over time. **D, E** Effect of Apratoxin A on expression of ribosome stalling (D) and protein degradation (E) reporters, related to Figure 2C and 2D, respectively. Graphs display mean +/- SD of GFP intensities at different time points (n = 3 independent treatments). **F** Effect of Apratoxin A (ApraA) on the glycosylation status of mCh-CFTR (24 h treatment). **G** Effect of Asterix and EMC5/6-KO on the expression of the mCh-CFTR protein degradation reporter (mean +/- SD of GFP intensity). p = unpaired t-test. Related to Figure 2E. **H** Validation of EMC-KO cell lines. Immunoblot for validation of EMC5, EMC6 and EMC5/6-dKO in HEK293 cells before (left) and after stable reporter integration (right). Bar chart shows protein degradation assay of a previously characterized EMC client, SOAT1 (Volkmar et al, 2019). **I** Terminal ribosome stalling analysis of K0 and CFTR reporters expressed in HEK293 WT, Asterix-KO, and EMC5/6-KO. mCh:GFP ratios (top) and GFP intensity (bottom) of n = 3 independent Dox treatments. P = unpaired t-test. Related to Figure 2F.

**Figure S6:**
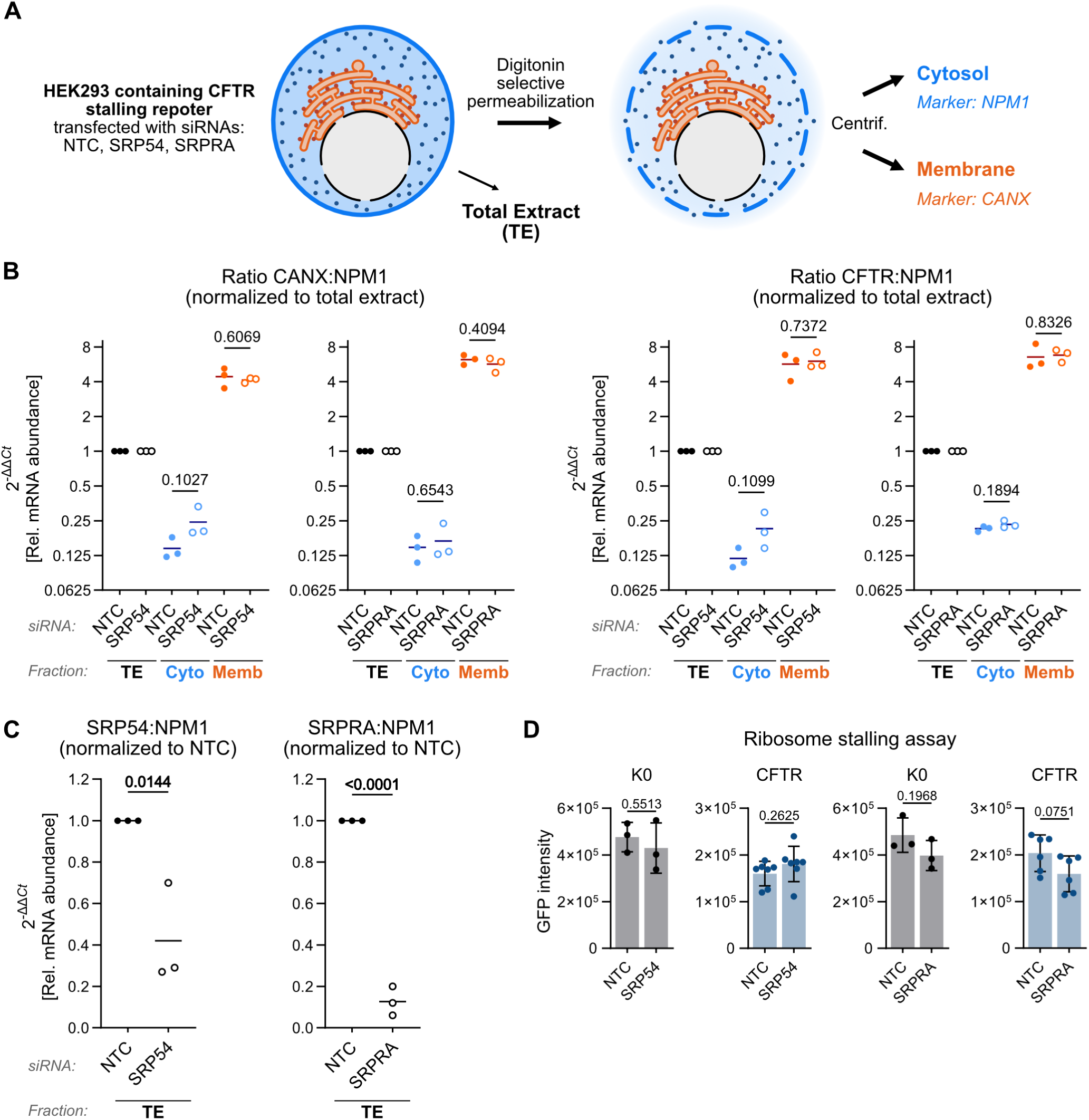
Digitonin fractionation of cytosolic and membrane-bound mRNAs in control and SRP pathway knockdown cells. **A** Schematic of fractionation experiment. Digitonin treatment selectively permeabilizes the plasma membrane, releasing cytosolic proteins and ribosomes while leaving endoplasmic reticulum membranes, membrane-associated proteins, ribosome-bound complexes, and portions of the cytoskeleton intact. Prior to permeabilization, two aliquots of the cells were collected for total RNA extraction and for flow cytometry analysis (Figure 3C). **B** Quantitative RT-PCR of total, cytosolic and membrane fractions, related to Figure 3C. Graphs depict the ratio of Calnexin (*CANX*, ER marker) and CFTR reporter mRNA levels in relation to *NPM1* transcript (cytosolic marker), normalized to the respective ratios in the total extract. For symmetry reasons, this ratiometric data is shown in Log 2 scale. p = unpaired t-test **C** Quantitative RT-PCR validation of SRP54 and SRPRA knockdown. p = unpaired t-test **D** Effect of SRP knockdown on the expression (mean +/- SD of GFP intensity) of K0 and CFTR stalling reporters. p = two-sided unpaired t-test. Related to Figure 3C.

**Figure S7:**
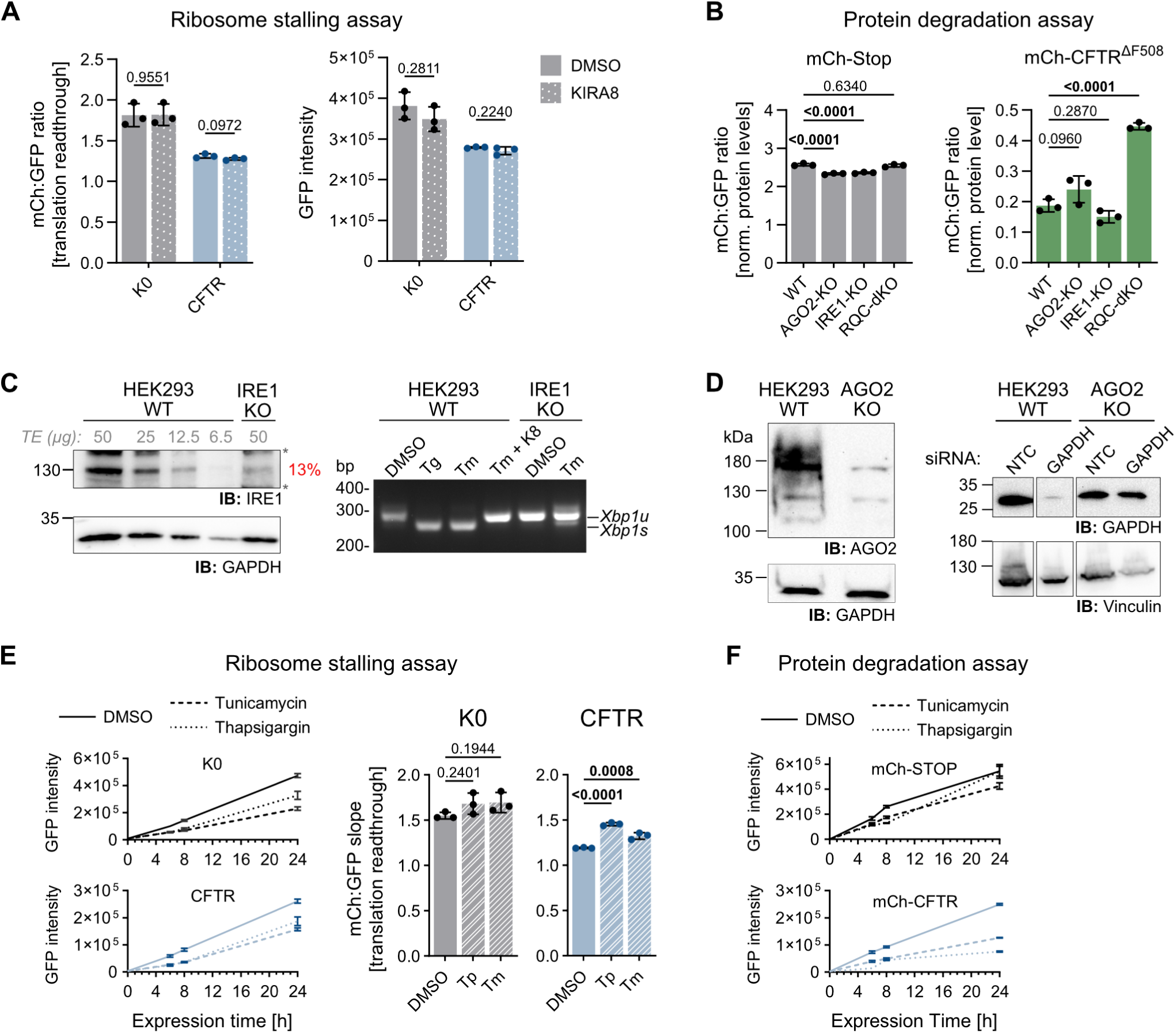
Influence of ER stress on abortive translation of CFTR. **A** Terminal ribosome stalling analysis of K0 and CFTR expressed in the presence of the IRE1 inhibitor KIRA8. Left: Mean +/- SD of mCh:GFP ratios from n = 3 independent treatments; same data as shown in Figure 4B, but not normalized to K0. Right: corresponding reporter expression levels (Mean +/- SD of GFP intensity). p = two-tailed unpaired t-tests. **B** Protein degradation assay of mCh-Stop and mCh-CFTR^ΔF508^ reporters transiently transfected in HEK293 WT, AGO2-KO, IRE1-KO and RQC-dKO cells. p = one-way ANOVA with Dunnet’s correction. **C** Validation of IRE1-KO in HEK293 cells. Left: Immunoblot analysis of IRE1 protein levels. Total protein extracts (TE) were loaded at different amounts. Estimated percentage of IRE1 signal in the polyclonal KO cell line is indicated in red. Right: Functional assay of IRE1 endonuclease activity. RT-PCR analysis of IRE1-mediated splicing of *XBP1* mRNA (s: spliced, u: unspliced) in the absence and presence of ER stressors Tunicamycin (Tm) or Thapsigargin (Tp) in HEK293 WT and IRE1-KO cells. The inhibitory effect of KIRA8 (K8) on Tm-induced IRE1 activity is also assessed in WT cells. **D** Validation of AGO2-KO in HEK293 cells. Left: Immunoblot analysis of AGO2 protein levels. Right: Functional assay of AGO2 activity. Immunoblot analysis of GAPDH protein levels in WT and AGO2-KO cells transfected with non-targeting control (NTC) and GAPDH*-*targeted siRNAs. **E** Terminal ribosome stalling analysis of K0 and CFTR reporters expressed in the presence of proteotoxic ER stressors Tm or Tp. Left: effect of ER stressors on reporter expression (mean +/- SD of GFP intensity) compared to DMSO. Right: bar graphs display mean +/- SD of mCh:GFP slopes from n = 3 independent treatments; same data shown in Figure 4C, but not normalized to K0. p = one-way ANOVA with Dunnet’s correction. **F** Effect of ER stressors on the expression (mean +/- SD of GFP intensity) of mCh-CFTR and mCh-Stop protein degradation reporters compared to DMSO, n = 3 independent treatments. Related to Figure 4D.

**Figure S8:**
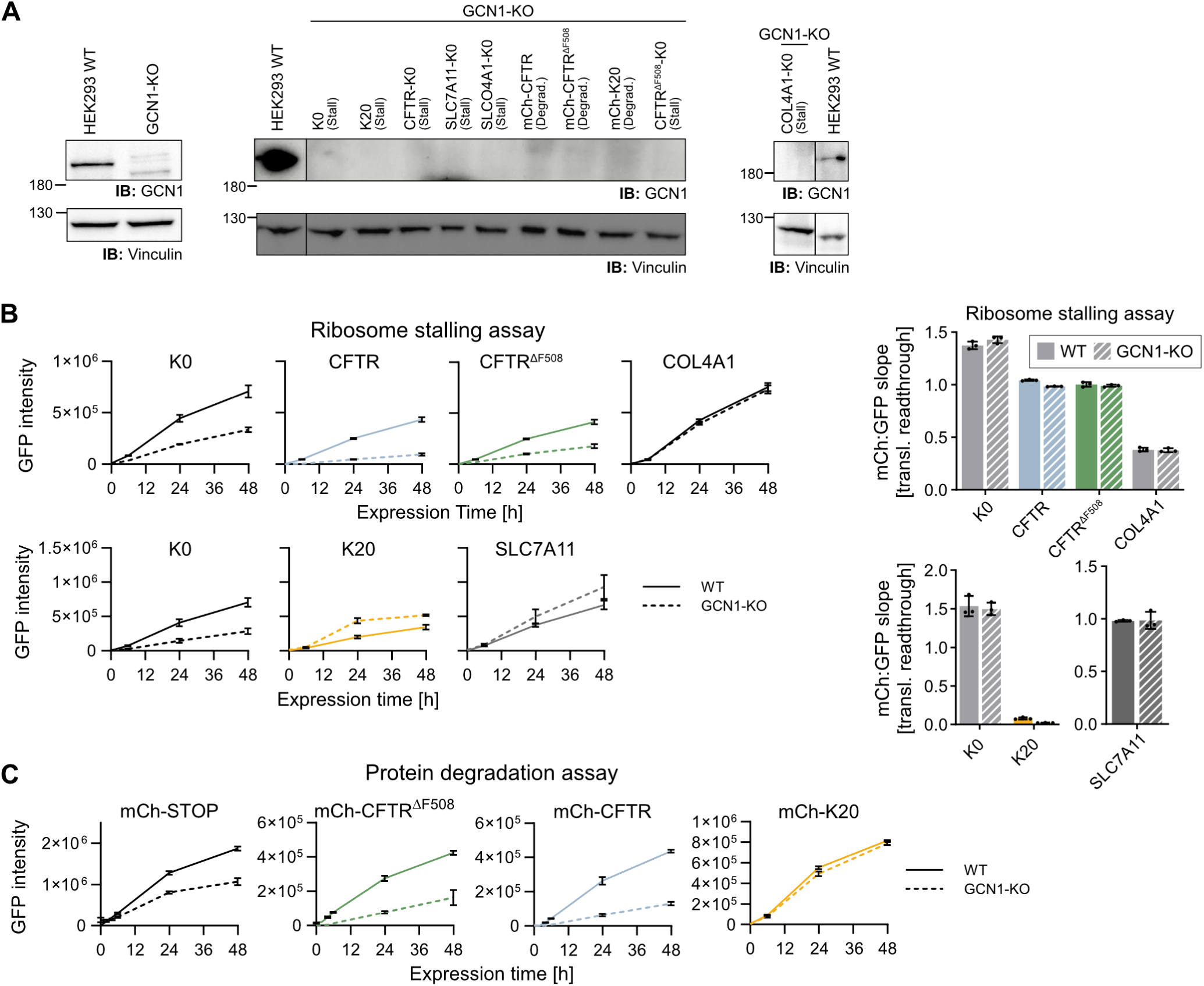
Influence of codon-optimality and collision sensor GCN1 on translation abortion. **A** Immunoblot analysis for validation of GCN1-KO in HEK293 cells before (left) and after stable reporter integration (middle and right). **B, C** Effect of GCN1-KO on expression of Dox-controlled genome-integrated ribosome stalling (B) and protein degradation (C) reporters. Graphs display mean +/- SD of GFP intensity from n = 3 independent Dox time course measurements of polyclonal reporter cell lines. Right: bar graphs display mean +/- SD of mCh:GFP slopes of ribosome stalling assay; same data as shown in Figure 5A, but not normalized to K0.

**Figure S9:**
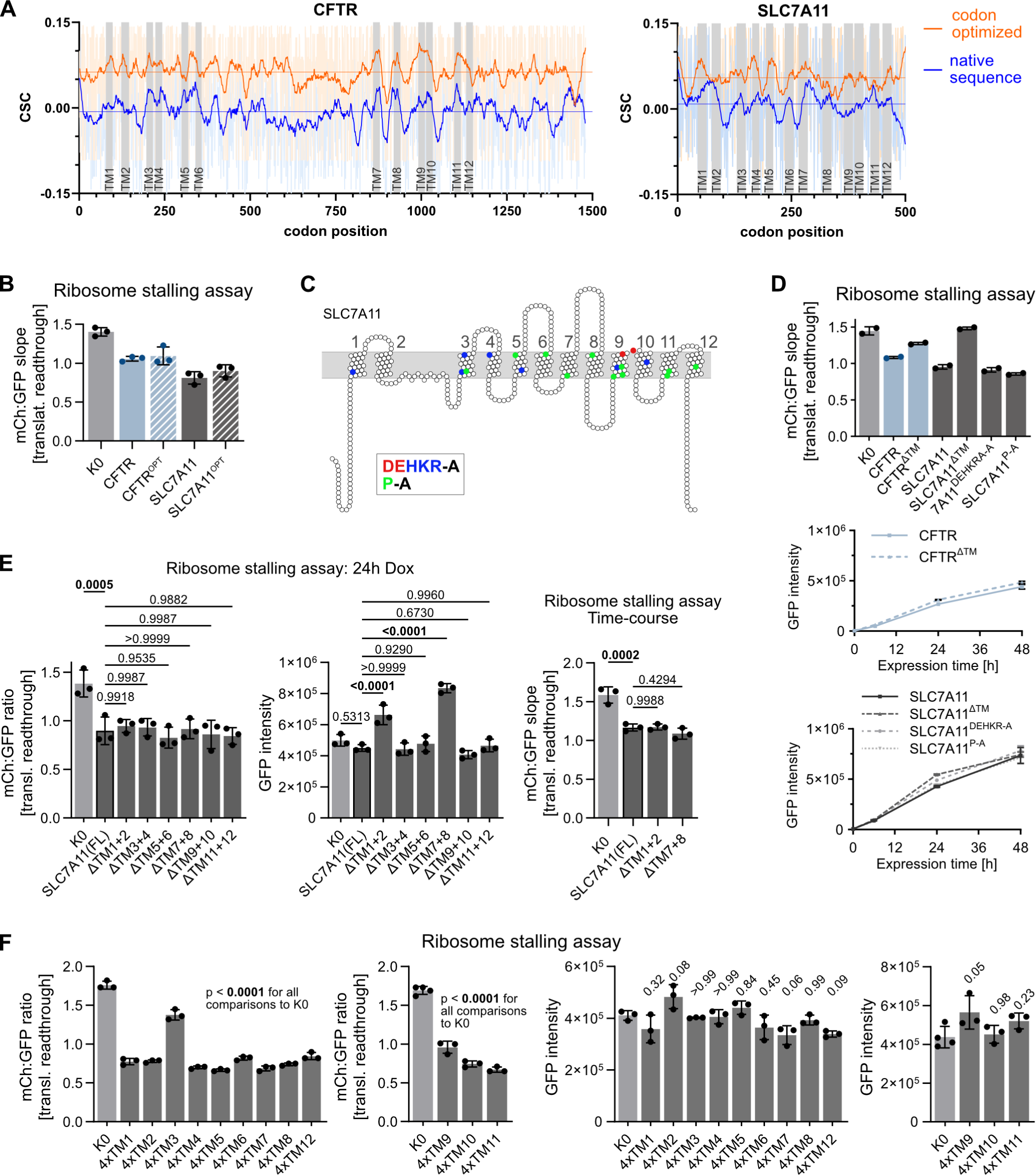
Analysis of the sequence determinants of terminal ribosome stalling of CFTR and SLC7A11. **A** Codon-stability coefficient (CSC) calculated for HEK293 wild-type cells (Müller *et al*., 2023)are used as proxy for codon-optimality. Thick lines represent smoothed CSCs (2^nd^ order polynomial, 20 neighbours on each side) for codons along the native (blue) or codon-optimized (orange) coding sequences of CFTR or SLC7A11. Gray shading indicates codons encoding for TM segments. **B** Effect of codon optimization on the CFTR and SLC7A11 terminal ribosome stalling reporters. Mean +/- SD of mCh:GFP slopes; same data as shown in Figure 6A, but not normalized to K0. **C** Cartoon illustrations of the SLC7A11 protein, indicating the position of the mutated amino acid residues of the SLC7A11^DEHKRA-A^ (charged residues - blue and red) and SLC7A11^P-A^ (proline residues - green) ribosome stalling reporters. Protein representations are drawn to scale of protein length, according to protein topology and protein feature annotation derived from UniProt with the help of the Protter interactive protein feature visualization tool (Omasits *et al*., 2014). **D** Terminal ribosome stalling analysis of CFTR and SLC7A11 TM mutants. Segments encoding for TMs were removed from the coding sequence (ΔTM). Charged residues (DEHKRA-A) or prolines (P-A) located within TM segments were mutated to alanine. Bar graph displays mean +/- SD of mCh:GFP slopes from n = 2 independent time course analyses of each polyclonal reporter cell line; same data as shown in Figure 6B, but not normalized to K0. Bottom: corresponding reporter expression (Mean +/- SD of GFP intensity) upon Dox induction over time. **E** Terminal ribosomal stalling analysis of SLC7A11 TM pair deletions. Left: Mean +/- SD of mCh:GFP ratios from n = 3 independent measurements (24 h Dox induction); same data as shown in Figure 6C, but not normalized to K0. p = one-way ANOVA with Dunnet’s correction, comparison to SLC7A11 full-length (FL). Center: corresponding reporter expression (Mean +/- SD GFP intensity). p = one-way ANOVA with Dunnet’s correction, comparison to SLC7A11 full-length (FL). Right: The SLC7A11 ΔTM1+2 and ΔTM7+8 reporters showed significantly increased reporter expression, and therefore their level of terminal ribosomal stalling was additionally measured in time course experiments. Mean +/- SD of mCh:GFP slope from n = 3 independent time course experiments. p = one-way ANOVA with Dunnet’s correction, comparison to SLC7A11 (FL). **F** Terminal ribosomal stalling analysis of K0+4xTM reporters. Top: Mean +/- SD of mCh:GFP ratios from n = 3 independent measurements (24 h Dox induction); same data as shown in Figure 6D, but not normalized to K0. p = one-way ANOVA with Dunnet’s correction, comparison to K0. Bottom: corresponding reporter expression (Mean +/- SD GFP intensity). p = one-way ANOVA with Dunnet’s correction, comparison to K0. The twelve reporters were measured in two sets of experiments, statistical comparison is done in relation to the K0 reference measured in the same replicate.

**Figure S10.**
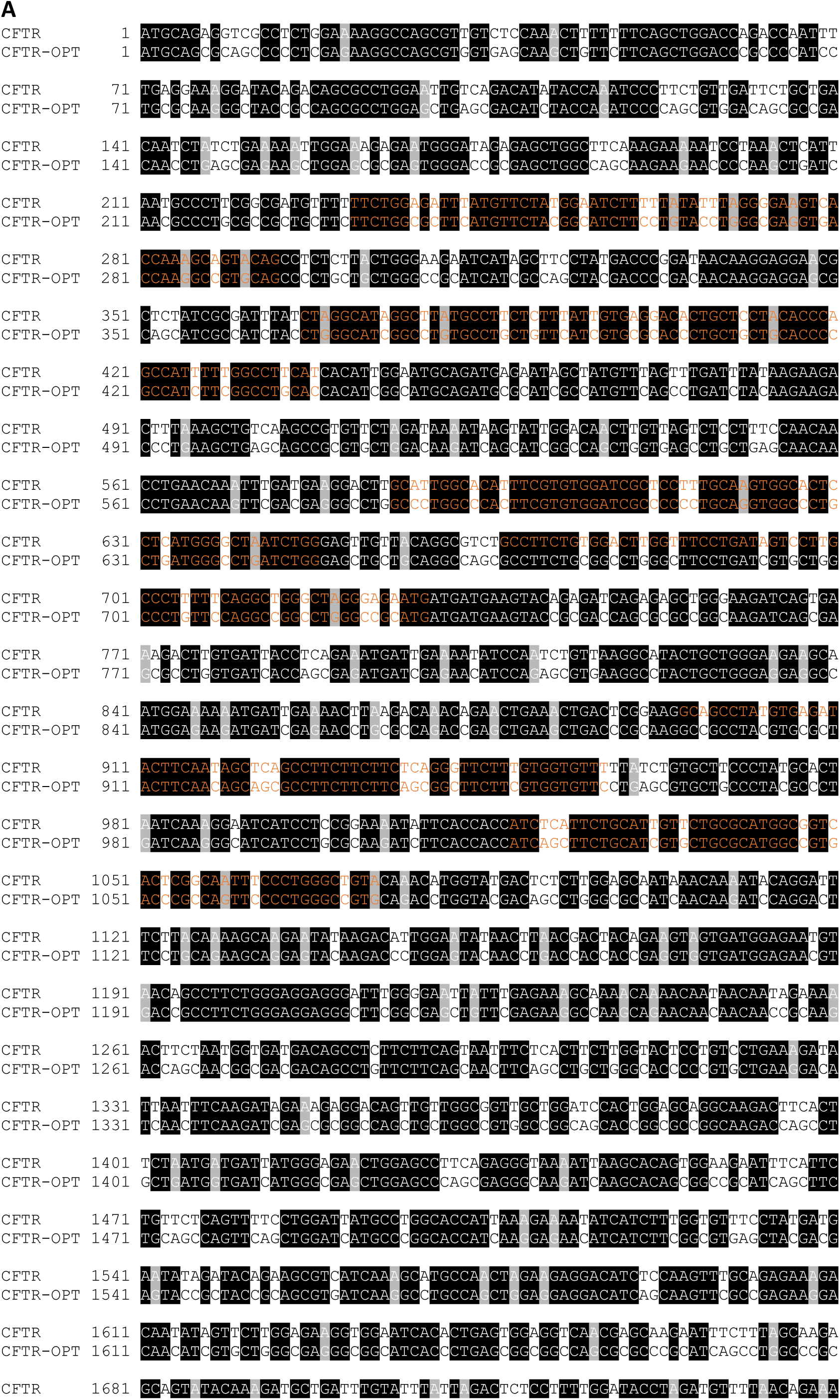
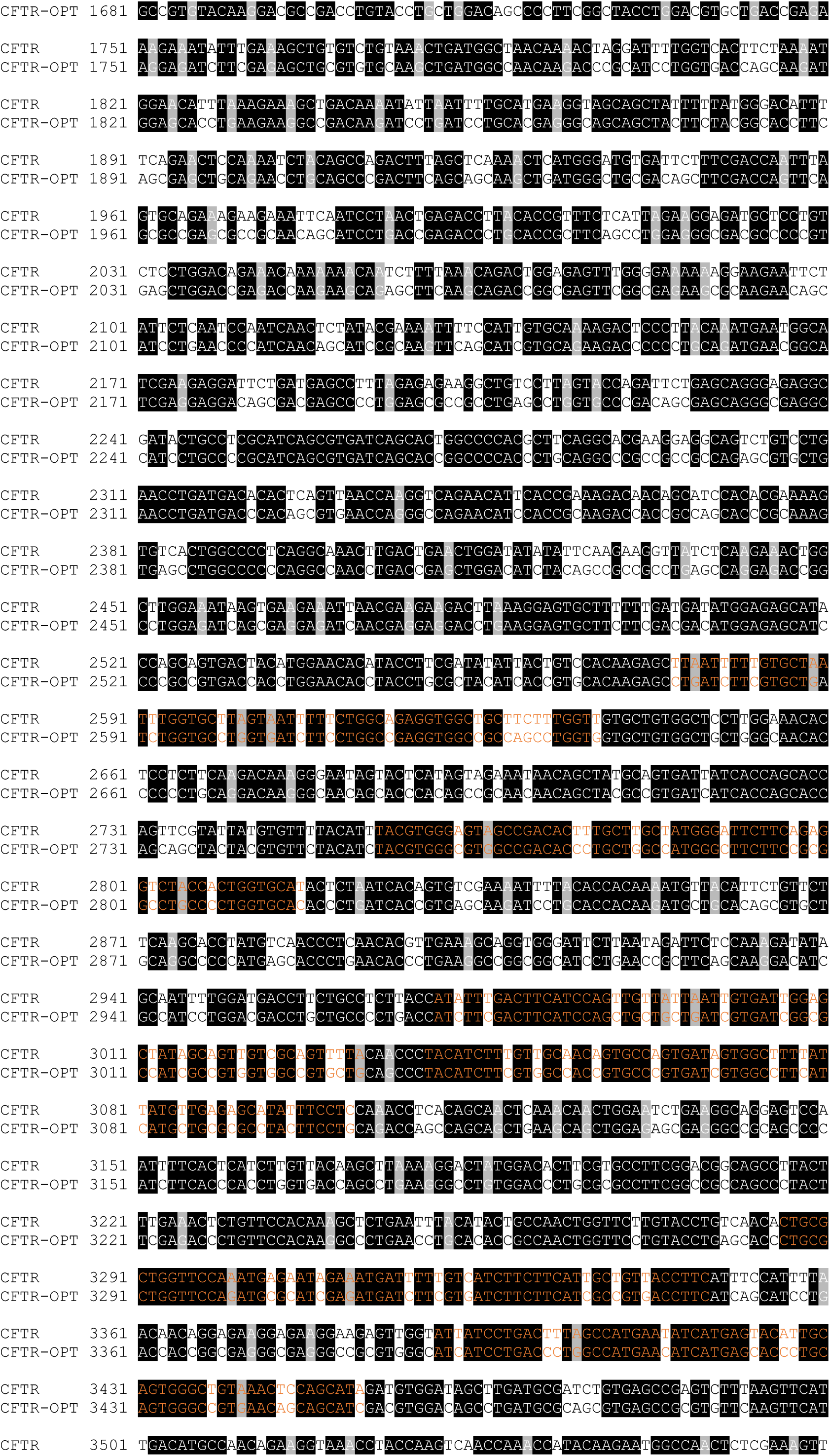
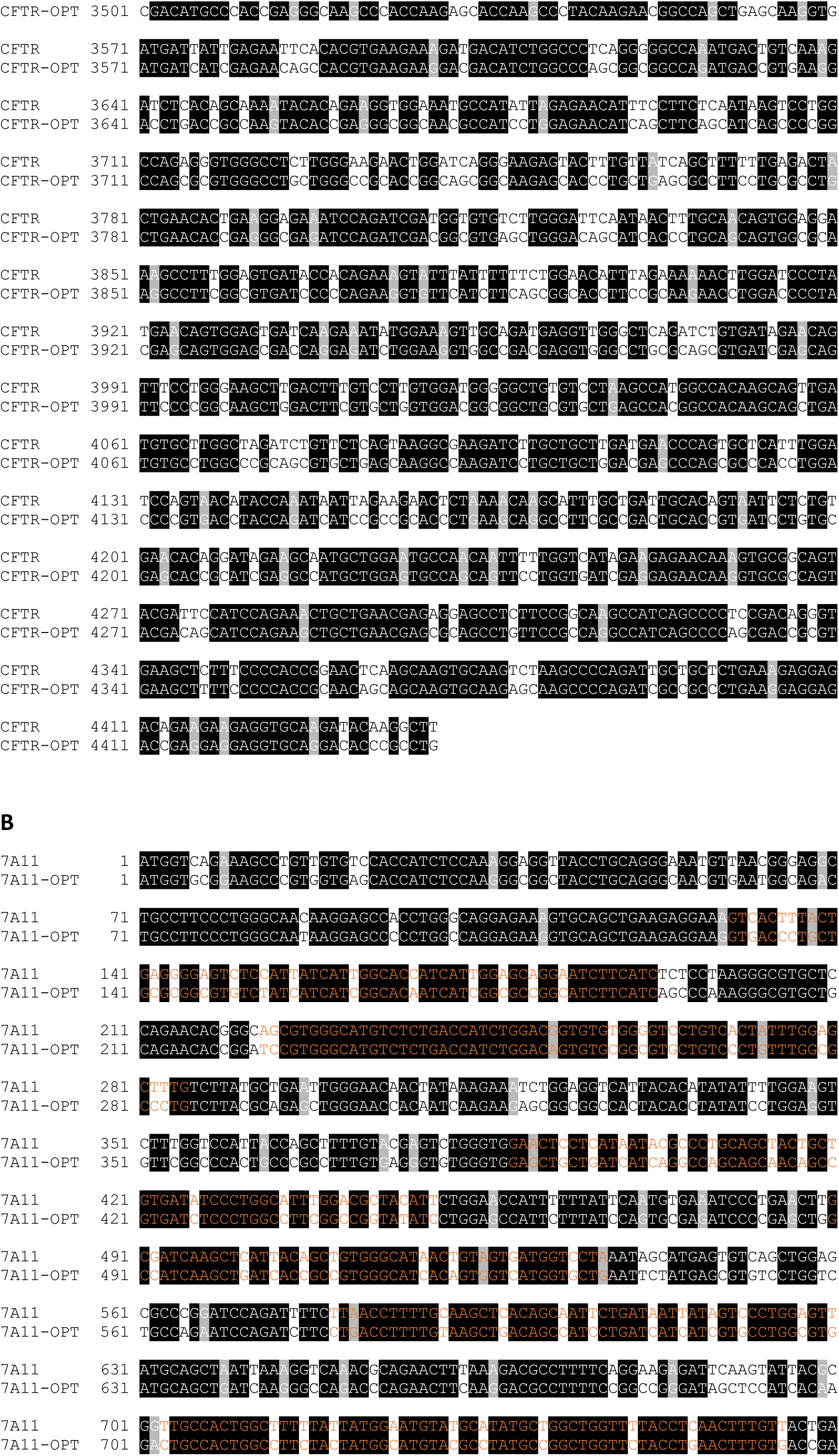
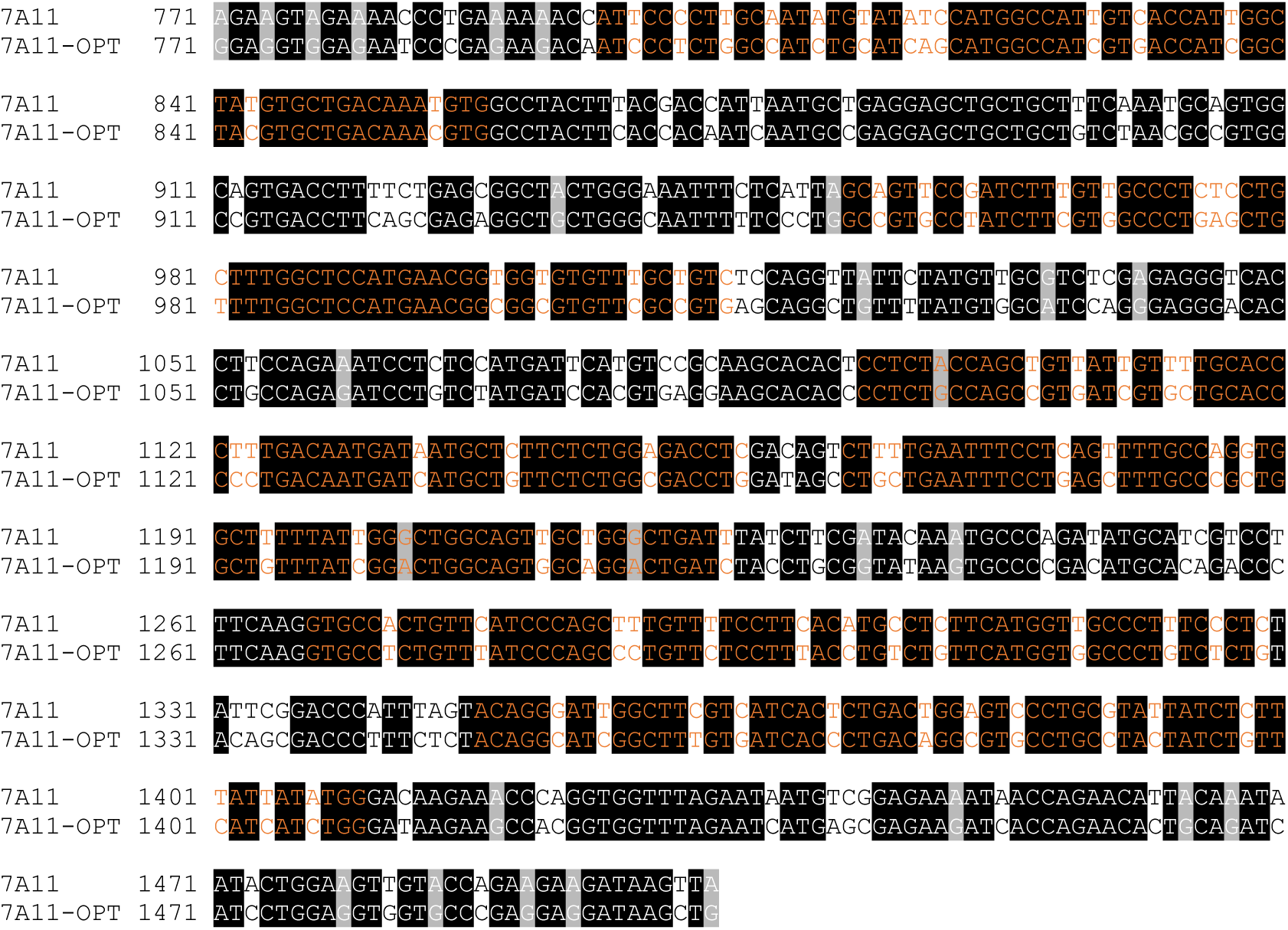
Sequence alignment of native and codon-optimized CFTR (**A**) and SLC7A11 (**B**). Segments encoding for TM regions are marked in orange.

